# The SPIRE1 actin nucleator coordinates actin/myosin functions in the regulation of mitochondrial motility

**DOI:** 10.1101/2020.06.19.161109

**Authors:** Felix Straub, Tobias Welz, Hannah Alberico, Rafael Oliveira Brandão, Anna Huber, Annette Samol-Wolf, Cord Brakebusch, Dori Woods, Martin Kollmar, Javier Martin-Gonzalez, Eugen Kerkhoff

## Abstract

Subcellular localisation of mitochondria provides a spatial and temporal organisation for cellular energy demands. Long-range mitochondrial transport is mediated by microtubule tracks and associated dynein and kinesin motor proteins. The actin cytoskeleton has a more versatile role and provides transport, tethering, and anchoring functions. SPIRE actin nucleators organise actin filament networks at vesicle membranes, which serve as tracks for myosin 5 motor protein-driven transport processes. Following alternative splicing, SPIRE1 is targeted to mitochondria. In analogy to vesicular SPIRE functions, we have analysed whether SPIRE1 regulates mitochondrial motility. By tracking mitochondria of living fibroblast cells from *SPIRE1* mutant mice and splice-variant specific mitochondrial SPIRE1 knockout mice, we determined that the loss of SPIRE1 function increased mitochondrial motility. The *SPIRE1* mutant phenotype was reversed by transient overexpression of mitochondrial SPIRE1, which almost completely inhibited motility. Conserved myosin 5 and formin interaction motifs contributed to this inhibition. Consistently, mitochondrial SPIRE1 targeted myosin 5 motors and formin actin filament generators to mitochondria. Our results indicate that SPIRE1 organises an actin/myosin network at mitochondria, which opposes mitochondrial motility.

**Summary statement:** The mitochondrial SPIRE1 protein targets myosin 5 motor proteins and formin actin-filament nucleators/elongators towards mitochondria and negatively regulates mitochondrial motility.

## Introduction

Animal cells have developed a sophisticated mitochondrial transport system in order to maintain energy homeostasis throughout the cell. As a major cellular energy source, mitochondria synthesise adenosine triphosphate (ATP) through oxidative phosphorylation. The mitochondria are actively distributed within cells, and are anchored at places with enhanced energetic demand (Chamberlain and Sheng, 2019). This is specifically important in cells with high energy demands and morphologically differentiated structure, such as neurons, where actin filament-dependent anchoring of mitochondria close to synapses is required to provide a local and stable energy supply for synaptic plasticity (Rangaraju et al., 2019).

Both transport and anchoring of mitochondria are mediated by elements of the cytoskeleton namely microtubules, actin filaments, and their associated motor proteins. Mitochondria are transported long-range along microtubule tracks in both directions by kinesin and dynein motors (Morris and Hollenbeck, 1995; Saxton and Hollenbeck, 2012). The motor proteins are targeted to the outer mitochondrial membrane by MIRO1/2 proteins of the RHO-GTPase family and the TRAK1 and TRAK2 adapter proteins (Fransson et al., 2006; Stowers et al., 2002; van Spronsen et al., 2013). MIRO1/2 also coordinates the association of the actin motor protein myosin 19 (MYO19) to mitochondria by direct interaction (Lopez-Domenech et al., 2018; Oeding et al., 2018). MYO19 is a plus end-directed actin motor protein. Transgenic expression of MYO19 was found to increase mitochondrial motility (Quintero et al., 2009). A major function of MYO19 is to ensure actin-dependent faithful mitochondria segregation during cell division (Rohn et al., 2014). In addition, MYO19 is also responsible for active mitochondrial transport into growing filopodia tips (Shneyer et al., 2017). The *Drosophila melanogaster* myosin 5 motor protein (Dm-MYO5) was also found to regulate mitochondrial motility. Inhibition of Dm-MYO5 by RNAi knockdown significantly enhanced the neuronal transport of mitochondria in both anterograde and retrograde directions (Pathak et al., 2010). These data indicate that MYO5 opposes microtubule-based mitochondrial movement, and suggest a role of MYO5 in actin-dependent mitochondrial docking. It is not yet known how *Drosophila* MYO5 is linked to mitochondria.

A potential mammalian MYO5 mitochondrial targeting factor is the SPIRE type actin nucleation factor 1 (SPIRE1), which was found to be localised at the outer mitochondrial membrane (Manor et al., 2015). SPIRE proteins interact directly via a conserved sequence motif in the central region (globular tail domain binding motif, GTBM) with the globular tail domain (GTD) of MYO5 motor proteins (Pylypenko et al., 2016). A short sequence motif (58 amino acids) encoded by an alternatively spliced exon mediates binding of SPIRE1 to mitochondria (Manor et al., 2015). The mitochondrial SPIRE1 isoform was originally termed SPIRE1C (Manor et al., 2015) and has been renamed here in accordance with its discoverer Uri Manor to mitoSPIRE1, because there an unrelated C-terminal *Drosophila* splice variant named SpireC previously existed (Rosales-Nieves et al., 2006).

SPIRE protein function is best studied in vesicle transport processes, where the mammalian SPIRE1 and SPIRE2 proteins organise actin filament/myosin 5 networks at the vesicle membranes, which transport RAB11 vesicles to the oocyte cortex in mice (Pfender et al., 2011; Schuh, 2011) and, in the case of melanocytes, mediate a centrifugal dispersion of RAB27 melanosomes (Alzahofi et al., 2020). Both vesicle transport processes are driven by MYO5 motor proteins (MYO5B in oocytes, MYO5A in melanocytes). In both cases the SPIRE proteins cooperate with FMN-subgroup formins (mammalian FMN1 and FMN2) to generate a vesicle-originated actin filament network. SPIRE/FMN cooperativity in actin filament generation is dependent on a direct interaction of the proteins, which is mediated by C-terminal FMN sequences (formin SPIRE interaction (FSI) motif), which interact with the SPIRE-KIND domain (Pechlivanis et al., 2009; Vizcarra et al., 2011; Zeth et al., 2011). For mitoSPIRE1, an additional formin interaction with inverted formin-2 (INF2, CAAX-isoform) has been described, and a cooperative function in actin filament generation at mitochondria/endoplasmic reticulum (ER) contact sites during mitochondrial division was proposed (Manor et al., 2015).

In accordance to MYO5/SPIRE/FMN cooperation in vesicle transport processes, herein we have addressed the question whether mitoSPIRE1 functions in the regulation of mitochondrial motility. Our data show that loss of mitoSPIRE1 function, similar to the loss of MYO5 function in *Drosophila* neurons, increases the motility of mitochondria. Consistently, we show that a gain of mitoSPIRE1 function stalls mitochondrial motility, depending on mitoSPIRE1 sequence motifs mediating the interactions with MYO5 and FMN-subgroup formins. We propose a model in which mitoSPIRE1 cooperates with FMN1 and class-5 myosin motor proteins in generating actin/myosin networks at mitochondria to anchor and position mitochondria in subcellular compartments.

## Results

### Origin and expression of mitoSPIRE1

The recent discovery of a mitochondrial SPIRE1 isoform (Manor et al., 2015) has opened the possibility of applying existing knowledge of SPIRE function in vesicle transport processes to the understanding of actin/myosin functions in mitochondrial dynamics. Currently, there is very little known about mitochondrial SPIRE1 function; therefore, we began by addressing basic knowledge on the origin and expression of mitoSPIRE1.

Phylogenetic analysis of SPIRE proteins of extant organisms traced the onset of SPIRE to the origin of the holozoa (ichthyosporea, choanoflagellates and metazoa)(Kollmar et al., 2019). Conversely, SPIRE proteins were not detected in plants and fungi. Genome duplications during chordate evolution gave rise to up to three *SPIRE* genes (*SPIRE1, SPIRE2, SPIRE3*) and alternatively spliced exons were introduced (Kollmar et al., 2019). The mammalian genomes encode two *SPIRE* genes, *SPIRE1* and *SPIRE2*. The *SPIRE2* gene encodes for only a single transcript, whereas the *SPIRE1* gene contains two alternatively spliced exons, exon 9 and exon 13 (Fig. 1A, green, sky blue), which give rise to three SPIRE1 isoforms, SPIRE1, SPIRE1-E9 and mitoSPIRE1, which contains the exon 13 encoded sequence (Fig. 1A; Fig. 2A). Gene expression analysis failed to detect a transcript that has both alternatively spliced exons (Kollmar et la., 2019). All SPIRE proteins encode an N-terminal kinase non catalytic C-lobe domain (KIND), which interacts with FMN-subgroup formins, a cluster of monomeric globular actin (G-actin) binding Wiskott-Aldrich homolgy 2 (WH2) domains, a myosin 5 globular tail domain binding motif (GTBM), a RAB interaction domain (Alzahofi et al., 2020) consisting of the SPIRE-box, a modified FYVE zinc finger domain (mFYVE), and conserved C-terminal flanking sequences (Kollmar et al., 2019)(Welz and Kerkhoff, 2019)(Fig. 1B). The function of the *SPIRE1* exon 9-encoded sequences is not known. The *SPIRE1* exon 13-encoded sequences are integrated in between the GTBM and the SPIRE-box and mediate the interaction of the mitoSPIRE1 protein with the outer mitochondrial membrane (Manor et al., 2015)(Fig. 1A, B; Fig. 2A, C, D; Fig. 3). Transient expression of AcGFP1-tagged vesicular and mitochondrial SPIRE1 proteins in human HeLa cells performed here revealed an exclusive localisation of the mitoSPIRE1 protein to mitochondria, and a complete elimination of vesicular SPIRE1 spots from mitochondria (Fig. 2D). The exon 13 encoded sequences are sufficient to target the AcGFP1 protein towards mitochondria (Fig. 2D). AcGFP1 alone did not display mitochondrial nor vesicular localisation.

**Fig. 1.**
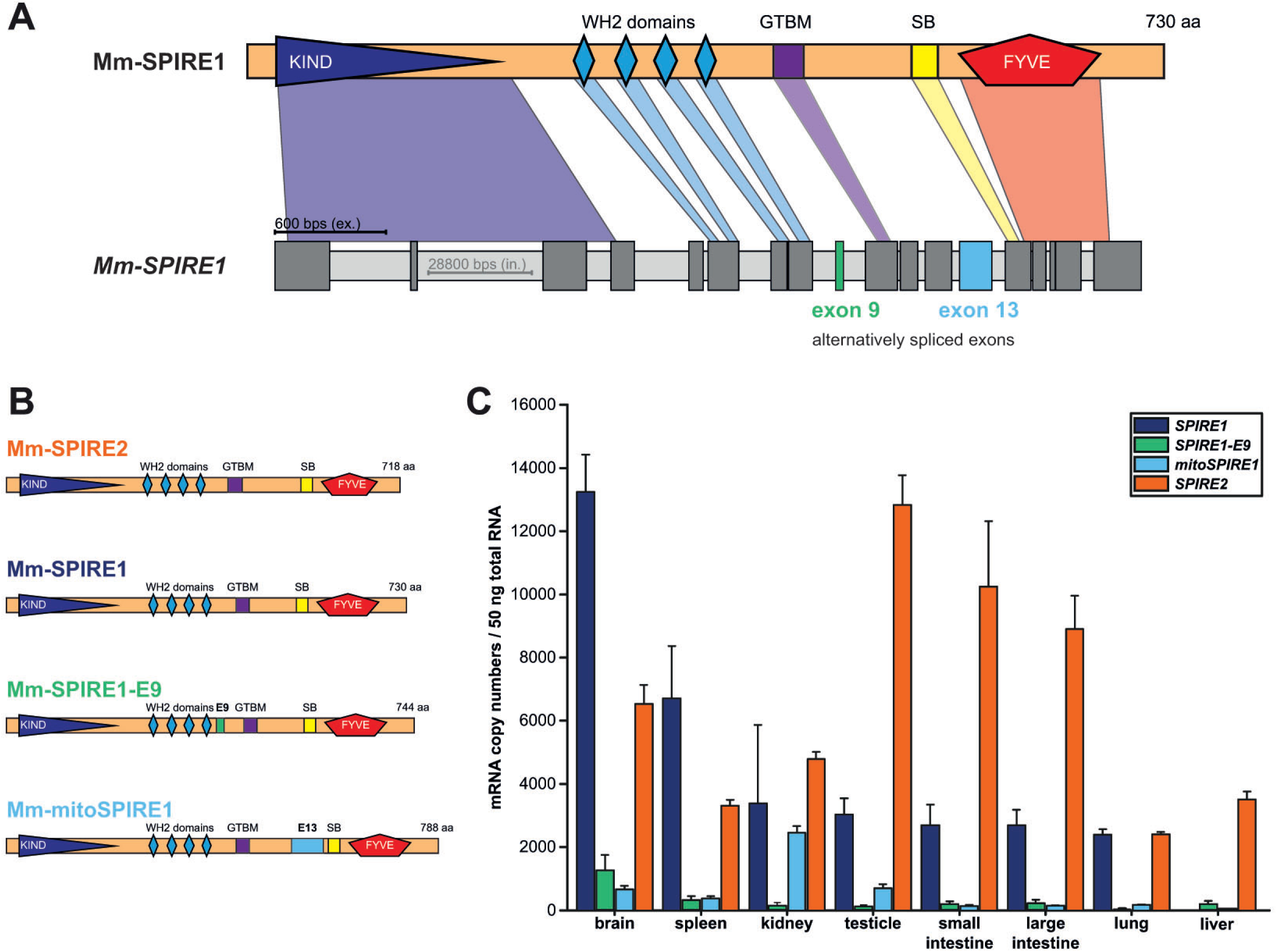
Structure and expression of mitoSPIRE1. The two mammalian *SPIRE* genes encode four different SPIRE proteins, which are highly similar but show different expression patterns. **(A)** Exon/intron structure of the mouse *SPIRE1* gene and domain composition of the corresponding SPIRE1 protein are shown. **(B)** Schematic overview on mouse SPIRE proteins and their respective domain organizations. Conserved domains between SPIRE proteins are highlighted and indicated. The *SPIRE1* gene encodes for three different proteins, SPIRE1, SPIRE1-E9 and mitoSPIRE1. SPIRE1-E9 contains differentially included exon 9, a unique 14 amino acid spanning sequence. The alternatively spliced exon 13, which encodes a sequence of 58 amino acids, is part of the mitoSPIRE1 protein. The *SPIRE2* gene encodes for one protein, SPIRE2. *KIND*, kinase non-catalytic C-lobe domain; *WH2*, Wiskott-Aldrich-Syndrome protein homology 2; *E9*, exon 9; *GTBM*, globular tail domain binding motif; *E13*, exon 13; *SB*, SPIRE-box; *FYVE*, FYVE-type zinc-finger; *aa*, amino acids; SPIRE1 accession-number: gi|149321426|ref|NC_000084.5|NC_000084 Mus musculus chromosome 18; SPIRE2 accession-number: gi|149361523|ref|NC_000074.5|NC_000074 Mus musculus chromosome 8. **(C)** *SPIRE* gene expression in various mouse tissues. Absolute *SPIRE* mRNA copy numbers were determined by quantitative real time PCR employing cDNA preparations from total RNA isolations and are presented in a bar diagram. Each bar represents mean mRNA copy numbers of three experimental repeats. *Error bars* represent SEM. n = 3 experimental repeats.

**Fig. 2.**
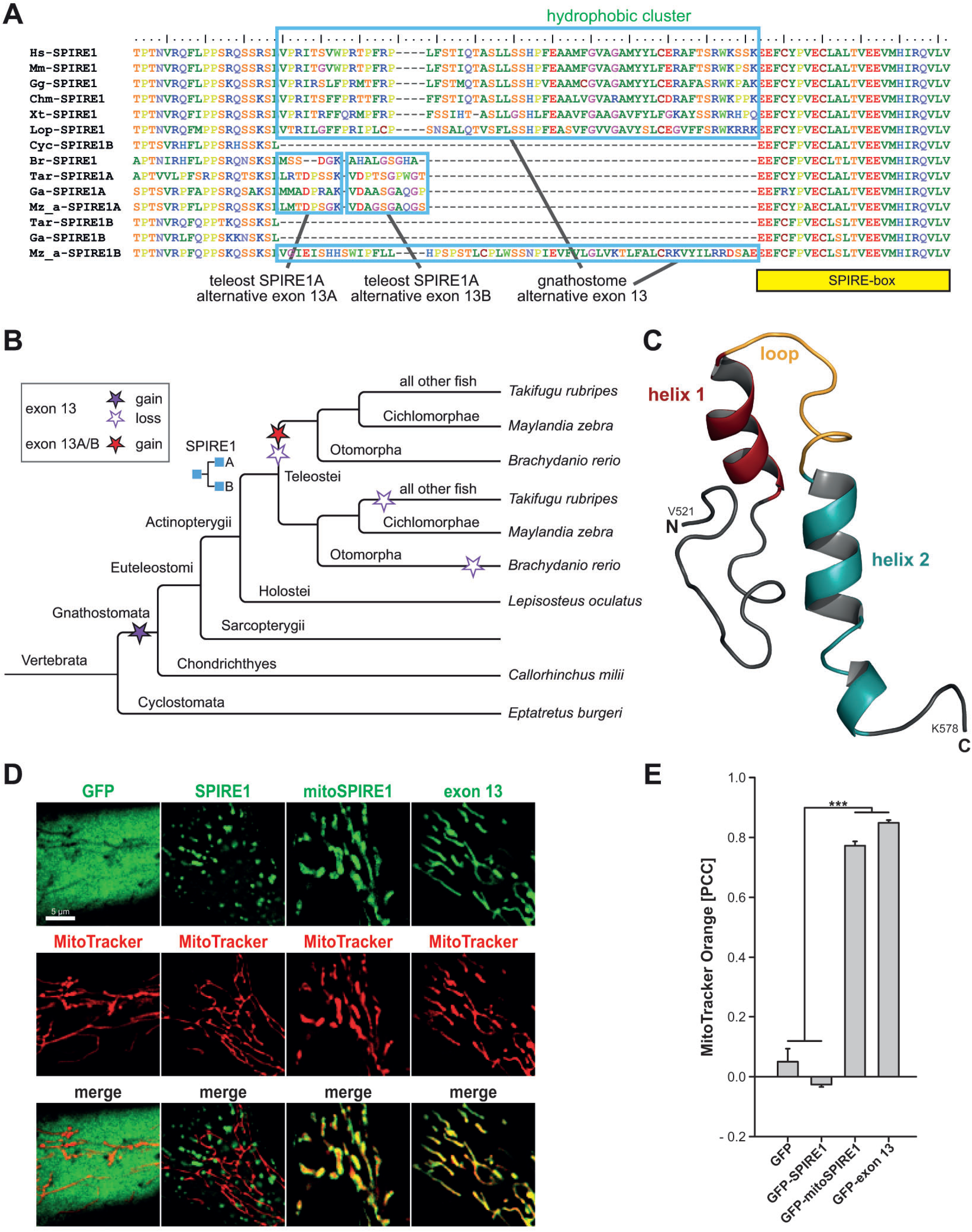
Evolution and predicted protein structure of *SPIRE1* exon 13 encoded sequences. **(A)** Multiple protein sequence alignment of selected SPIRE1 homologs showing the regions of and around the alternatively spliced exon 13. Species abbreviations: Br = *Brachydanio rerio*; Chm = *Chelonia midas*; Cyc = *Cyprinus carpio*; Ga = *Gasterosteus aculeatus*; Gg = *Gallus gallus*; Hs = *Homo sapiens*; Lop = *Lepisosteus oculatus*; Mm = *Mus musculus*; Mz_a = *Maylandia zebra;* Tar = *Takifugu rubripes*; Xt = *Xenopus tropicalis*. **(B)** Schematic phylogenetic tree of SPIRE1 homologs showing gain and loss of the alternatively spliced *SPIRE1* exon 13 during evolution. **(C)** Predicted secondary protein structure of the alternatively spliced mouse *SPIRE1* exon 13 (XM_006526204.3; amino acids 521 - 578) as was predicted by the PHYRE software (Kelley et al., 2015). Exon 13 is predicted to form two alpha helices (helix 1, *red* and helix 2, *petrol blue*) separated by a loop structure (*yellow*) allowing to form a hairpin structure for integration into the mitochondrial outer membrane. N-terminal (N) and C-terminal (C) end of the exon 13 are distinguished. Amino acid labelling is provided for better orientation. (D) Cytoplasmic localisation of mitochondrial and vesicular SPIRE1 proteins. AcGFP1 and AcGFP1-tagged SPIRE1,mitoSPIRE1, and the isolated exon 13 sequences have been transiently expressed in HeLa cells. Mitochondria have been visualised by MitoTracker orange staining. Fluorescence microscopy analysis showed co-localisation of AcGFP-mitoSPIRE1 and AcGFP-E13 with mitochondria. The AcGFP1-SPIRE1 had a vesiclular localisation, which was distinct from the mitochondria. AcGFP1 alone had an evewn cytoplasmic distribution.

**Fig. 3.**
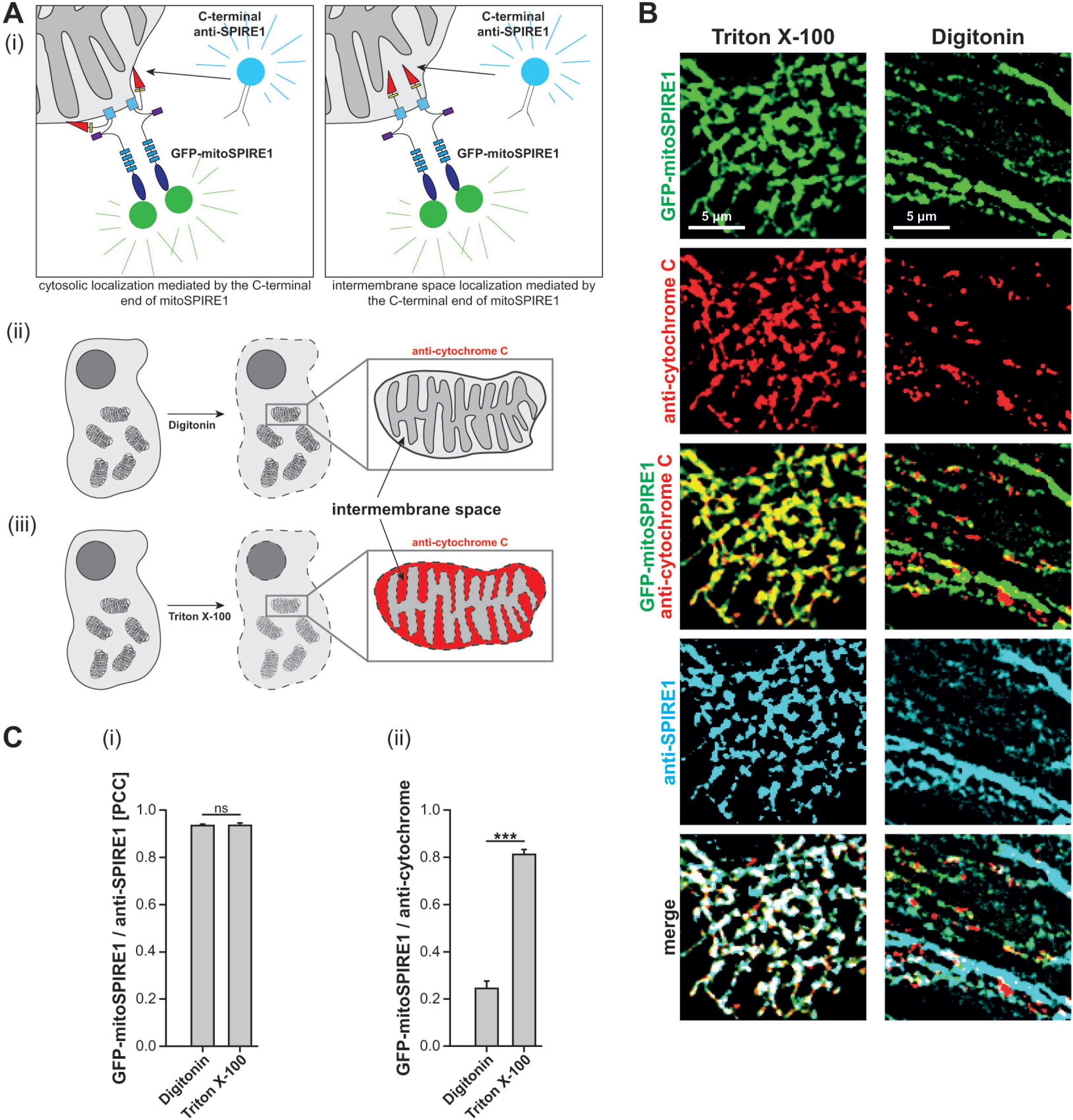
Interaction of mitoSPIRE1 with the outer mitochondria membrane. The presence of the C-terminal end of mitoSPIRE1 in the cytosol or within the mitochondrial intermembrane space was analysed. **(A)** Scheme of the experimental setup. Digitonin is used to permeabilize only the plasma membrane leaving the outer mitochondrial membrane intact, while Triton X-100 is used to permeabilize both, the plasma membrane and organellar membranes. Therefore, only Triton X-100 treatment allows entry of antibodies into the mitochondrial intermembrane space (IMS) to reveal specific protein staining. **(i)** A C-terminus specific SPIRE1 antibody (C-terminal anti-SPIRE1, SA-2133, *cyan*) is used to indicate the C-terminal end of transiently expressed GFP tagged mitoSPIRE1 (mitoSPIRE1, N-terminal GFP tag, *green*). **(ii)** Antibody staining of mitochondria located cytochrome C (located in the IMS) was used as a control showing that the outer mitochondrial membrane is still intact after Digitonin (indicated by light grey color of the IMS). **(iii)** Triton X-100 treated cells also show permeabilized mitochondrial membranes (indicated by dashed lines membrane) allowing antibody staining of cytochrome C proteins (indicated by *red* color of IMS). **(B)** GFP tagged mitoSPIRE1 proteins (AcGFP1; GFP-mitoSPIRE1, *green*) are transiently expressed in HeLa cells. Deconvoluted images show the localization of transiently expressed GFP-mitoSPIRE1 proteins (*green*) at mitochondrial membranes, an antibody staining of the C-terminal end of mitoSPIRE1 (*cyan*) and an IMS antibody staining of cytochrome C (*red*). Fixed cells were permeabilized for antibody staining by Triton X-100 or by the mild permeabilizer Digitonin. Comparison of merge pictures from GFP-mitoSPIRE1 (*green*) and cytochrome C (*red*) shows overlapping localization of cytochrome C and GFP-mitoSPIRE1 in Triton X-100 but not in Digitonin treated cells. At least 10 cells were recorded for each condition and one representative cell is shown here. *Scale bar* represents 5 μm. **(C)** The colocalization of GFP tagged mitoSPIRE1 with cytochrome C and C-terminal SPIRE1 signals, respectively, as described in (B) was quantified by determining its Pearson’s correlation coefficient (PCC) and is shown in a bar diagram. Each bar represents the mean PCC value for at least six cells analyzed. **(i)** The PCC for colocalization of GFP-mitoSPIRE1 and C-terminal SPIRE1 signal does not differ between Digitonin and Triton X-100 permeabilized cells. Mann-Whitney-U-test has been performed as post-hoc analysis: *ns* = not significant. *Error bars* represent SEM. **(ii)** Digitonin permeabilized cells show a significantly lower colocalization rate of cytochrome C (*red*) and GFP-mitoSPIRE1 (*green*) compared to cells permeabilized with Triton X-100. T-test has been performed as post-hoc analysis: *** = p < 0.001. *Error bars* represent SEM.

Differentially included *SPIRE1* exons are not present in the single *Eptatretus burgeri* (inshore hagfish) *SPIRE* homolog (Fig. 2B). The *Callorhinchchus milii* (Australian ghostshark, cartilaginous fishes) *SPIRE1* already contains the around 60 amino acids long, differentially included exon before the SPIRE-box motif indicating its origin in the ancient gnathostomian (jawed vertebrates) SPIRE homolog. The differentially included exon 13 is conserved between tetrapods, the single *Lepisosteus oculatus* (spotted gar) *SPIRE1*, and the *Cichlomorphae* (rayfinned fish, *SPIRE1B*). The alternative exon 13 has been lost in all other fish *SPIRE1B genes*, and has been substituted in *SPIRE1A* subfamily members by a set of two very short exons (Fig. 2A, B), of which each can be differentially included independently. The function of these short differentially included *SPIRE1A* exons is unknown.

Our phylogenetic data show that the mitochondrial SPIRE1 function originated after the two whole genome duplications that occurred early in vertebrate evolution and is conserved since with the exception of a subset of bony fish, where the sequences are either lost or substituted by short sequence motifs of unknown function (Fig. 2A, B).

In order to evaluate which mouse tissues express the mitoSPIRE1, we have performed an absolute quantification of *SPIRE1* mRNA by means of real-time quantitative polymerase chain reaction (qPCR; Fig. 1C). Specific primer pairs for each of the *SPIRE1* splice variants and *SPIRE2* have been employed to differentiate between the individual isoforms. Cloned cDNAs encoding for the specific mouse SPIRE isoforms were used as standards for absolute quantification of mRNA copy numbers in 50 ng of total RNA from the analysed tissues. In general, *SPIRE* expression is very low as compared to housekeeping genes. We confirmed the previously reported high expression of *SPIRE1* in brain, and the preferential expression of *SPIRE2* in tissues of the digestive tract (Pleiser et al., 2010; Schumacher et al., 2004)(Fig. 1C). In most cases, the expression of the isoforms containing the alternatively spliced *SPIRE1* exon 9 and exon 13 was very low as compared to both the *SPIRE1* isoform without alternative exons, as well as the *SPIRE2* isoform. Interestingly, we found very high expression of the exon 13-encoding *mitoSPIRE1* isoform in kidney (Fig. 1C), which might reflect the important role of mitochondrial function in energetically demanding kidney filtration processes.

### Interaction of mitoSPIRE1 with the outer mitochondrial membrane

The exon 13-encoded sequences insert a 58 amino acid spanning motif in between the myosin 5 and RAB binding motifs of mitoSPIRE1, which is sufficient for to target SPIRE1 to mitochondria (Manor et al., 2015)(Figs. 1, 2, 3). In order to evaluate the mode of interaction with the outer mitochondrial membrane, we have performed a structure prediction (PHYRE, protein homology/analogy recognition engine V 2.0 (Kelley et al., 2015) of the exon 13-encoded sequence. The proposed helix-loop-helix structure and the highly hydrophobic character of the motif (Figs. 2A, C) suggest that mitoSPIRE1 could bind the outer mitochondrial membrane by a helical loop insertion into the outer mitochondrial membrane lipid bilayer. If this were the case, however, the RAB-binding sequences (SPIRE-box, mFYVE and flanking sequences) would be outside of the mitochondria, which would be in contradiction to published data, which suggested that C-terminal mitoSPIRE1 protein sequences penetrate the outer mitochondrial membrane, and are localised in the mitochondrial intermembrane space (Manor et al., 2015)(Fig. 3A).

To address the localisation of the C-terminal end of mitoSPIRE1, we have performed immunofluorescence microscopy experiments, employing an anti-SPIRE1 antibody, which recognises a peptide at the C-terminal end of mitoSPIRE1 (anti-SPIRE1-CT; (Schumacher et al., 2004). The detergent Digitonin, which permeabilises the plasma membrane for antibody staining at low concentrations, was incapable of permeabilizing the outer mitochondrial membrane under these conditions (Vercesi et al., 1991)(Fig. 3A). In contrast, the detergent Triton X-100 permeabilises both membranes for antibody staining. Co-immunostaining of HeLa cells transiently expressing an AcGFP1-tagged mitoSPIRE1 protein (GFP-mitoSPIRE1) with antibodies against the mitoSPIRE1 C-terminus and the mitochondrial intermembrane space marker cytochrome C resulted in a co-immunostaining of GFP-mitoSPIRE1 and cytochrome C only in the case of the permeabilization with Triton-X 100, which allows the penetration of antibodies into the mitochondria (Figs. 3A, B, C). In Digitonin-permeabilised cells, where no antibody penetration into the mitochondria occurs, we detected GFP-mitoSPIRE1 with anti-SPIRE1-CT antibodies. Here, immunofluorescence perfectly overlapped with the auto-fluorescence of the AcGFP1-tagged mitoSPIRE1 protein (Figs. 3A, B, C). Conversely, there was only a minor overlap between cytochrome C staining and the AcGFP1/anti-SPIRE1-CT staining (Figs. 3B, C), indicating that the Digitonin did not enable anti-cytochrome C antibody penetration into the mitochondria. This signifies that the C-terminus of mitoSPIRE1 is localised outside of the mitochondrial outer membrane, and is thus assessible for antibody staining under Digitonin permeabilization conditions.

In summary, these results suggest that the exon 13-encoded sequence motif anchors mitoSPIRE1 to mitochondria by the insertion of a hydrophobic helical loop into the outer mitochondrial membrane, indicating that the C-terminal end of mitoSPIRE1 is located in the cytosol and not in the mitochondrial intermembrane space.

### Generation of a splice-variant specific mitoSPIRE1 knockout mouse model

Research on SPIRE function in vesicle transport processes has been complicated in the past by redundant SPIRE1 and SPIRE2 functions (Pfender et al., 2011)(Alzahofi et al., 2020). The mitochondrial SPIRE1 function, however, cannot be compensated by SPIRE2 since SPIRE2 does not localise to mitochondria (Kollmar et al., 2019). A specific knockout of the mitoSPIRE1 splice variant by a deletion of exon 13 should, therefore, completely impair mitochondrial SPIRE1 function. In order to address the cell biological functions of mitoSPIRE1, we decided to generate mitoSPIRE1 knockout cells and compare mitochondrial dynamics of the knockout cells with those of wild-type cells. We have followed the strategy to generate a mitoSPIRE1 knockout mouse model as a source for primary mitoSPIRE1 knockout fibroblasts and future studies on mitoSPIRE1 function in kidney homeostasis and neuronal plasticity. The knockout was achieved by CRISPR/Cas9-mediated targeted deletion of the alternatively spliced *SPIRE1* exon 13 in mouse embryonic stem cells (Fig. 4A). Subsequently, mitoSPIRE1 knockout mice were derived from the knockout stem cells. The deletion of exon 13 was confirmed by a PCR analysis of chromosomal DNA employing the use of primers upstream and downstream of the deletion site (Fig. 4A, B).

**Fig. 4.**
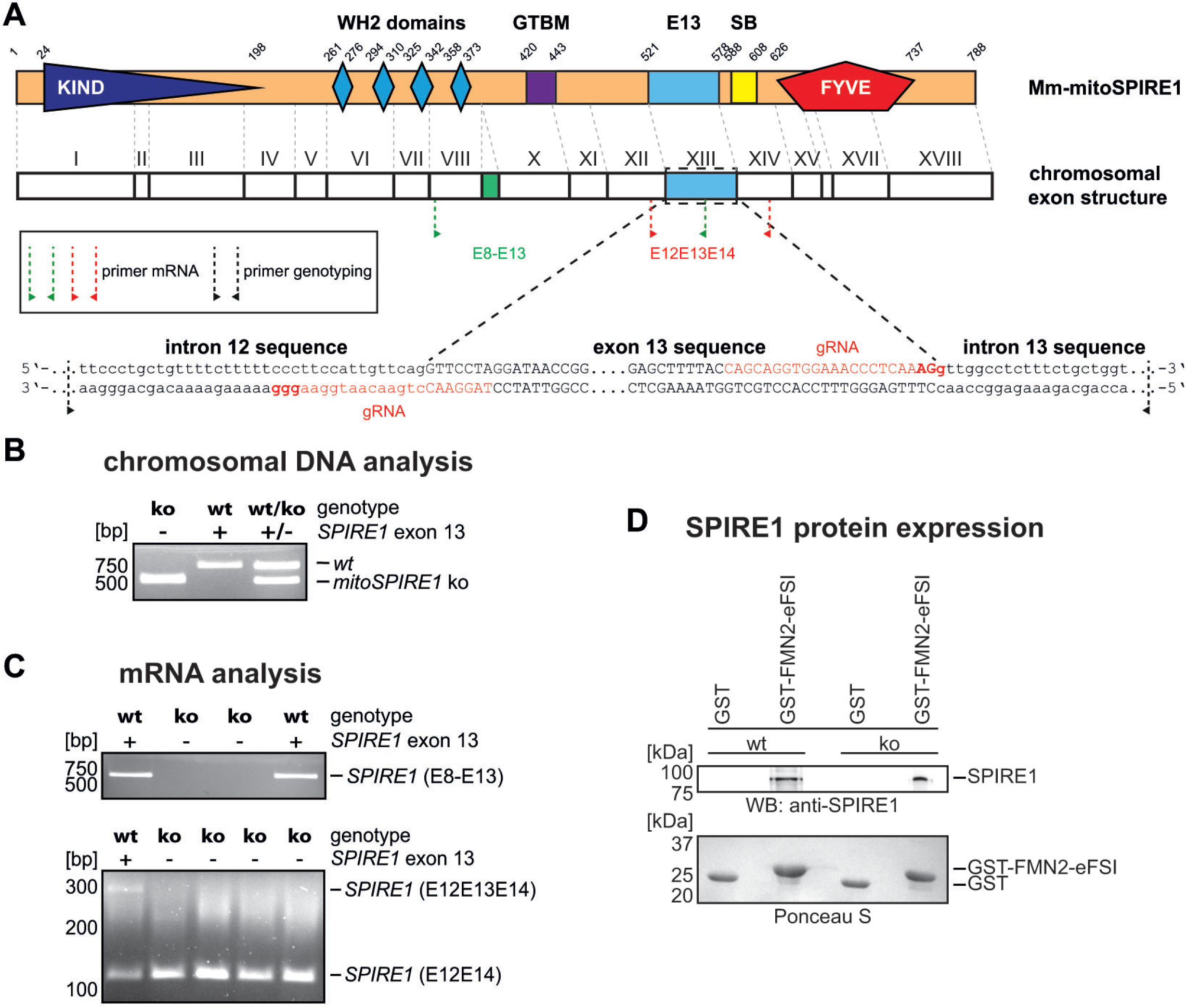
Generation of the mitoSPIRE1 knockout mouse. **(A)** Schematic representation of the mouse mitoSPIRE1 protein domains and the respective chromosomal exon structure. Numbers on the protein structure indicate amino acid boundaries. The genomic DNA sequence of exon 13 is shown (capital letters) as are the surrounding intron sequences (lowercase letters). Guide RNAs (gRNA) for CRISPR / Cas9 mediated gene editing are highlighted in *red*. Primer pairs used to verify excision of exon 13 sequences at genomic DNA and mRNA levels are illustrated as dotted arrows (DNA primer = *black*, mRNA primer = *green* and *red*). *KIND*, kinase non-catalytic C-lobe domain; *WH2*, Wiskott-Aldrich-Syndrome protein homology 2; *GTBM*, globular tail domain binding motif; *E13*, exon 13; *SB*, SPIRE-box; *FYVE*, FYVE-type zinc-finger. SPIRE1 accession-number: gi|149321426|ref|NC_000084.5|NC_000084 Mus musculus chromosome 18. **(B)** Proof for absence of *SPIRE1 exon 13* at genomic and mRNA levels but intact SPIRE1(-E9) protein expression in mitoSPIRE1 knockout mice. **(i)** Absence of genomic *SPIRE1 exon 13* in mitoSPIRE1 knockout mice was confirmed by PCR analysis using genotyping primer pairs (black dotted arrow, (A)). Genomic DNA of wild-type mice was used as control. **(ii)** RT-PCR analysis of cDNA preparations from primary embryonic fibroblasts (pMEFs) reveals absence of the alternatively spliced *exon 13* at mRNA levels in mitoSPIRE1 KO mice. Two different primer pairs were used (*red* and *green* dotted arrows as in (A) for PCR fragments *SPIRE1* (*E8-E13*) and *SPIRE1* (*E12E13E14*), respectively). **(iii)** SPIRE1 and SPIRE1-E9 proteins are still expressed in mitoSPIRE1 KO pMEFs as was confirmed by a GST-pulldown assay from pMEF cell lysates using purified GST-Mm-FMN2-eFSI (GST-FMN2-eFSI) and immunoblotting (anti-SPIRE1 antibodies, SA-2133). GST was used as control and Ponceau S staining shows equal amounts of GST tagged proteins. *WB*, Western blotting; *wt*, wild-type; *ko*; mitoSPIRE1 knockout.

As a cell culture system to study mitochondrial dynamics, primary mouse embryonic fibroblasts (pMEFs) were isolated from wild-type and knockout mouse embryos. Fibroblasts were chosen because they are specifically suitable for microscopic analysis of mitochondria dynamics due to their flat cell morphology. *SPIRE1* gene expression was analysed by reverse transcriptase PCR (RT-PCR) of mRNA isolated from mitoSPIRE1 knockout pMEFs and wild-type pMEFs. The RT-PCR experiments showed that the *SPIRE1* mRNA was still expressed and confirmed a specific loss of *SPIRE1*-exon 13 containing RNAs (Fig. 4C). Western blot analysis of wild-type and mitoSPIRE1 knockout fibroblasts employing an anti-SPIRE1 antibody, which recognises all SPIRE1 isoforms (anti-SPIRE1-CT (Schumacher et al., 2004)), confirmed the RT-PCR results in that the general expression of SPIRE1 is not influenced by the deletion of the exon 13 sequences (Fig. 4D). Since the expression of SPIRE1 proteins is very low, we had to enrich SPIRE1 proteins by GST-FMN2-eFSI pulldown to detect endogenous SPIRE1 in immunoblot experiments.

### Loss of mitoSPIRE1 function increases mitochondrial motility

Vesicular SPIRE proteins function in organising actin/myosin networks to drive vesicle transport processes towards the cell periphery (Pfender et al., 2011; Schuh, 2011)(Alzahofi et al., 2020). The nature of the transport pathways is determined by the interaction of SPIRE proteins with RAB GTPases, vesicle membranes, and MYO5 motor proteins (Pylypenko et al., 2016; Tittel et al., 2015)(Kollmar et al., 2019; Alzahofi et al., 2020). The *SPIRE1* exon 13-encoded sequences anchor mitoSPIRE1 to the mitochondrial membrane (Manor et al., 2015)(Fig. 3). Corresponding to the function of vesicular SPIRE proteins, we have addressed the question of whether mitoSPIRE1 influences mitochondrial motility.

To monitor the motility of mitochondria in living primary mouse fibroblasts we have stained cells with the fluorescent dye MitoTracker Orange, which is cell membrane permeable and specifically stains mitochondria. Live cell fluorescence microscopy was then employed to track the motility of mitochondria, which was later analysed with the Bitplane IMARIS particle tracking software module (Fig. 5, 7). To inhibit SPIRE1 function we have isolated primary fibroblasts from two different mouse models, the mitoSPIRE1 knockout mouse and the *SPIRE1* mutant mouse. The *SPIRE1* mutant mouse has a gene trap insertion in the *SPIRE1* gene, which abrogates SPIRE1 expression in between the KIND and WH2 domains (Pleiser et al., 2014). *SPIRE1* mutant mice do not express functional SPIRE1 proteins, and thereby lack vesicular and mitochondrial SPIRE1 functions. The motility of mitochondria in primary fibroblast cells from both mouse models was compared to that of primary fibroblasts isolated from wild-type mouse embryos. Our results show that the lack of mitoSPIRE1 function in mitoSPIRE1 knockout and in *SPIRE1* mutant primary fibroblasts significantly increases nearly all tested motility parameters. These parameters include the percentage of moving mitochondria, the average track length of a moving mitochondrion, the maximal track length, and the maximal velocity (Fig. 5A, B).

**Fig. 5.**
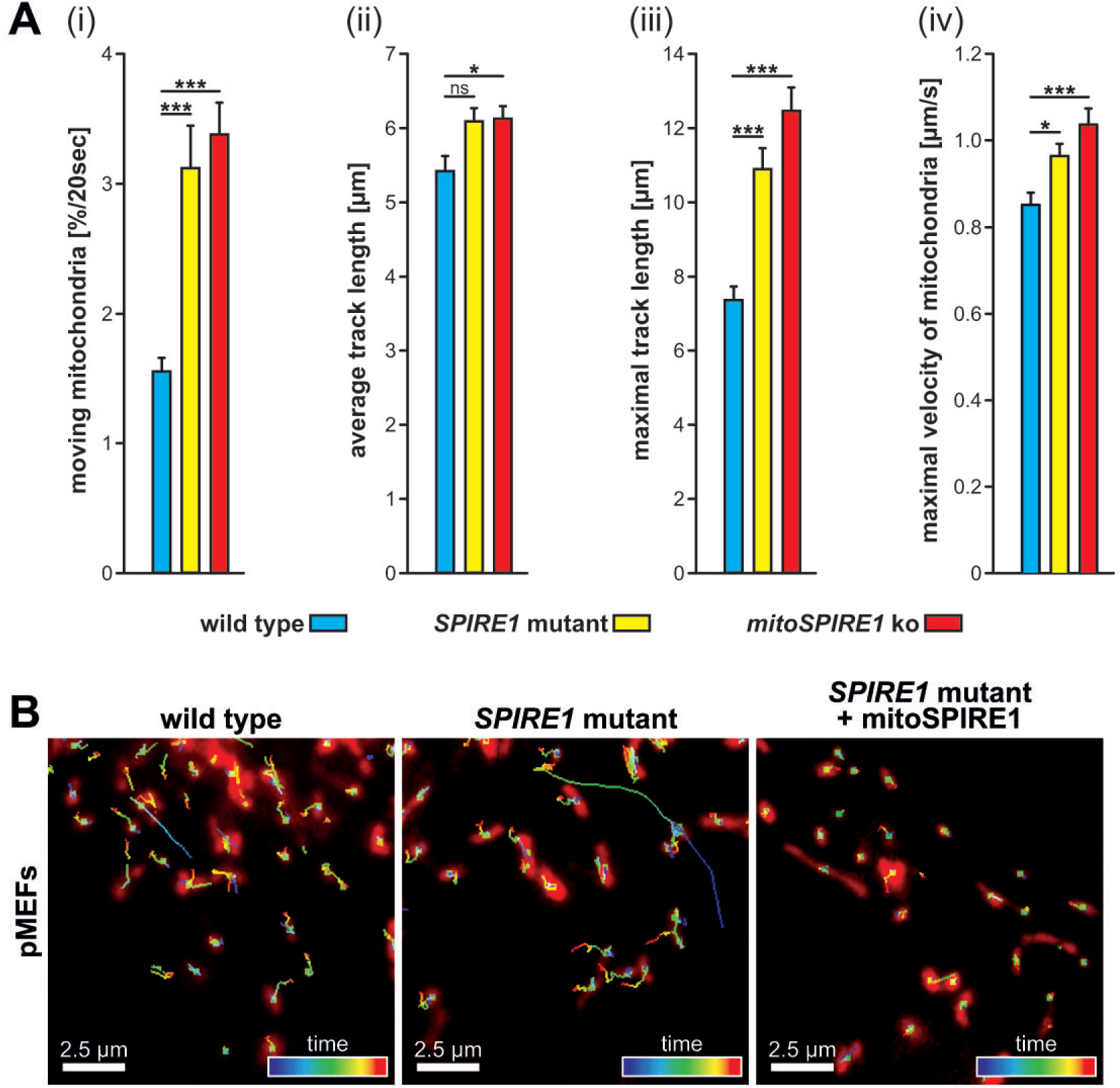
Loss of mitoSPIRE1 function increases mitochondrial motility. Motility of mitochondria was measured in a time range of 20 seconds in wild-type, *SPIRE1* mutant and mitoSPIRE1 knockout (mitoSPIRE1 KO) primary mouse embryonic fibroblasts (pMEFs) and migratory tracks were analyzed via IMARIS software. Motility of mitochondria was grouped according to the measured track length into three different classifications: *fixed, wiggle* and *moving* mitochondria. **(A)** The absence of mitoSPIRE1 increases the motility of mitochondria in pMEFs of *SPIRE1* mutant and mitoSPIRE1 KO mice compared to wild-type mice as shown in bar diagrams. In detail loss of mitoSPIRE1 significantly increased the number of moving mitochondria **(i)**, the average mitochondria migrational track length **(ii)**, the maximal migrational track length of moving mitochondria **(iii)** and the maximal transport velocity of moving mitochondria **(iv)**. Each bar represents mean values, *error bars* represent SEM. n > 39 cells per group. One-way ANOVA has been performed; for each parameter a significant difference between species was detected; proofed effect size [(i): η^2^ = 0.249; (ii): η^2^ = 0.054; (iii): η^2^ = 0.279; (iv): η^2^ = 0.112]; Tukey-Kramer test has been performed as post-hoc analysis: ns = not significant, * = p < 0.05, *** = p < 0.001. **(B)** Representative cells with MitoTracker Orange stained mitochondria including individual mitochondrial motility tracks for wild-type (left panel), *SPIRE1* mutant (middle panel) and *SPIRE1* mutant pMEFs overexpressing mitoSPIRE1 protein (right panel). Track color code represents timepoints of mitochondria movement. Mitochondria of *SPIRE1* mutant cells show a longer maximal track length compared to wild-type mitochondria which was reversed by overexpression of mitoSPIRE1.

These data indicate a general function of mitoSPIRE1 in negative regulation of mitochondrial motility. Our results are in contrast to SPIRE function in vesicle transport processes, where SPIRE-organised actin/myosin networks were shown to drive RAB27 and RAB11 vesicle transport (Pfender et al., 2011; Schuh, 2011)(Alzahofi et al., 2020).

**Table 1.**
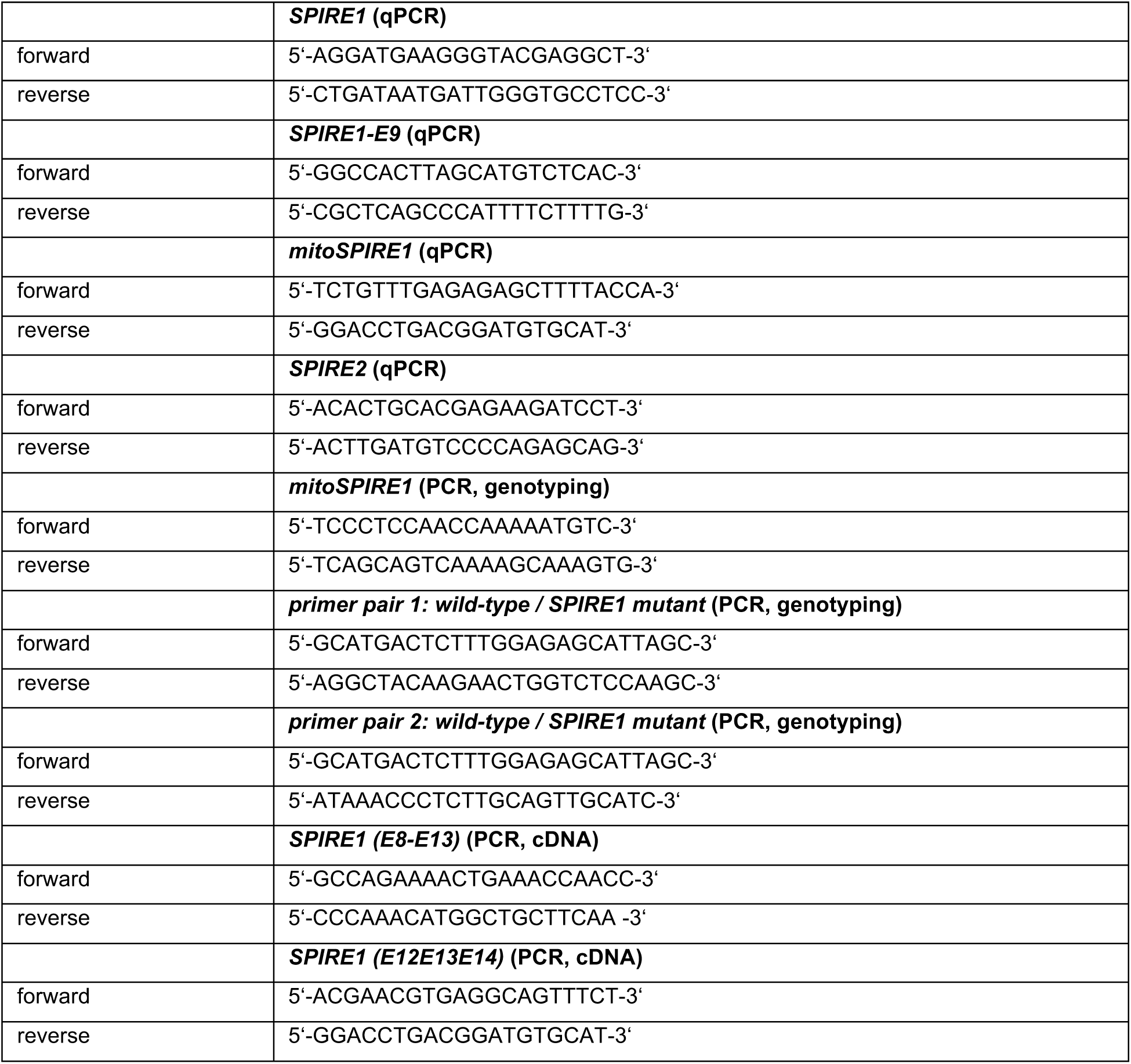
Specific primer pairs used for qualitative and quantitative PCRs and for genotyping.

**Table 2.**
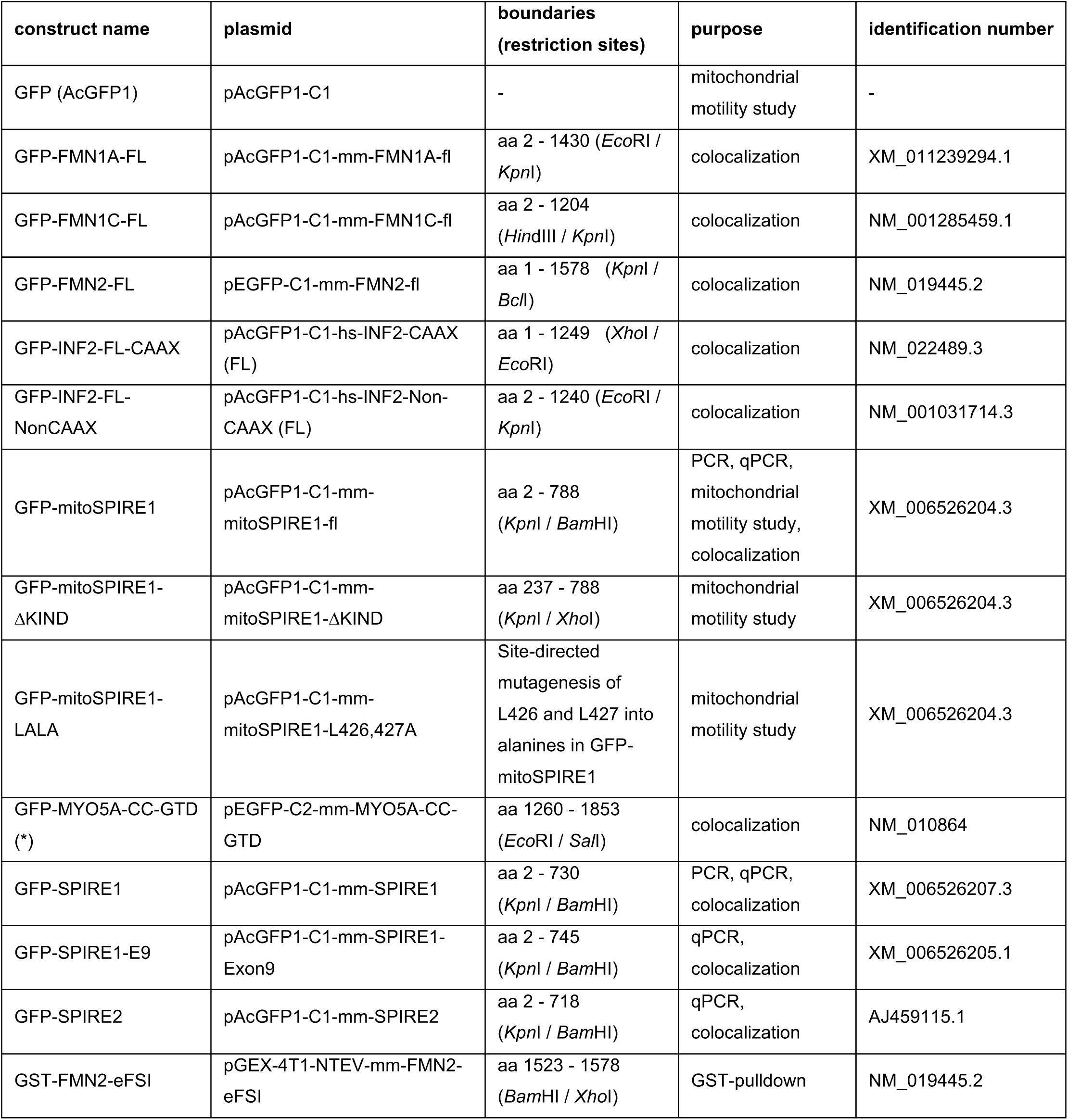
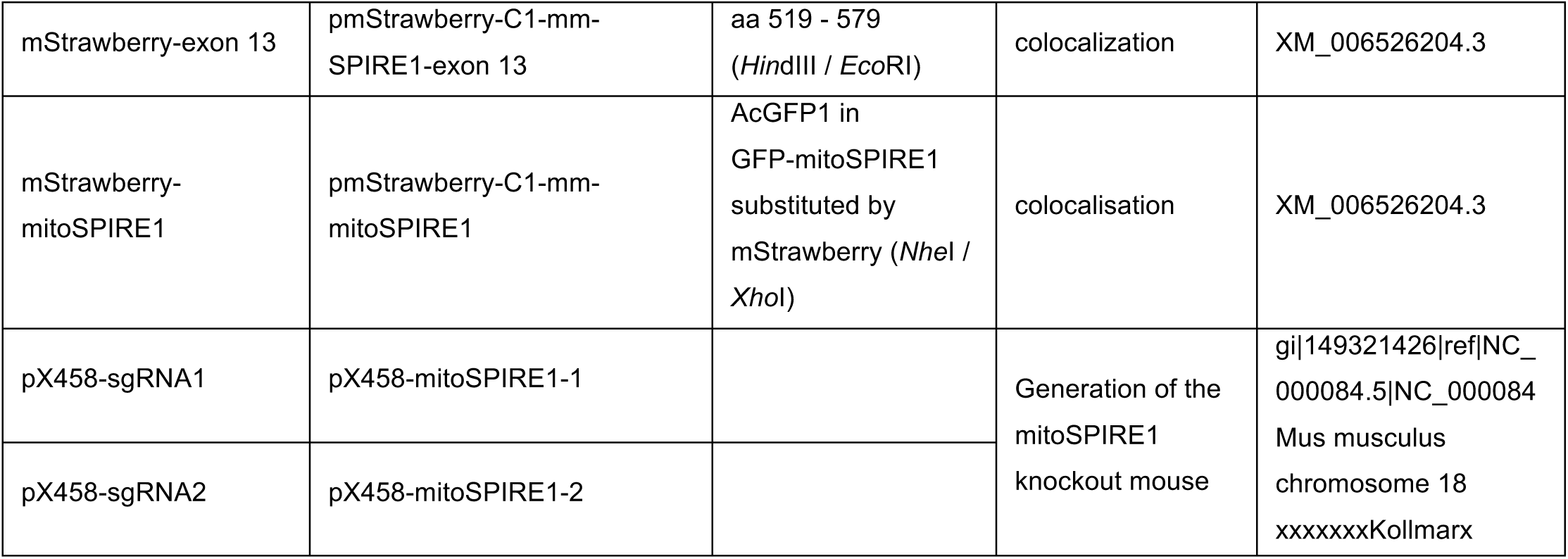
Overview on eukaryotic and prokaryotic expression vectors and respective cDNA accession numbers.

### mitoSPIRE1 targets myosin 5 function towards mitochondria

The inhibitory function of mitoSPIRE1 in vesicle transport processes very much resembles the phenotype of the myosin 5 knockout in fly neurons, where the loss of myosin 5 function also increased the motility of mitochondria (Pathak et al., 2010). SPIRE proteins can directly interact with class 5 myosin motor proteins via a conserved myosin 5 globular tail domain binding motif (GTBM) in the central part of the proteins (Pylypenko et al., 2016). The MYO5 binding motif is also encoded by the mitoSPIRE1 protein. We therefore asked whether mitoSPIRE1 acts as an adaptor for MYO5 motor proteins at the outer mitochondria membrane.

As a proof of principle, we have transiently co-expressed mStrawberry-tagged mitoSPIRE1 and eGFP-tagged C-terminal myosin 5A encoding the cargo binding coiled-coil (CC) region and the globular tail domain (GTD)(GFP-MYO5A-CC-GTD) in human HeLa cervix carcinoma cells (Fig. 6A, B). The localisation of the proteins was then analysed by fluorescence microscopy. We found a strong co-localisation of mitoSPIRE1 and the C-terminal MYO5A-CC-GTD protein (Fig. 6B). Co-expression of GFP-MYO5-CC-GTD with the mitochondria localised isolated *SPIRE1* exon 13 encoded sequences (mStrawberry-exon 13) revealed a separated vesicular and mitochondrial localisation of the proteins (Fig. 6B). eGFP alone did not colocalise with mStrawberry-mitoSPIRE1, nor did mStrawberry colocalise with GFP-MYO5A-CC-GTD (data not shown).

**Fig. 6.**
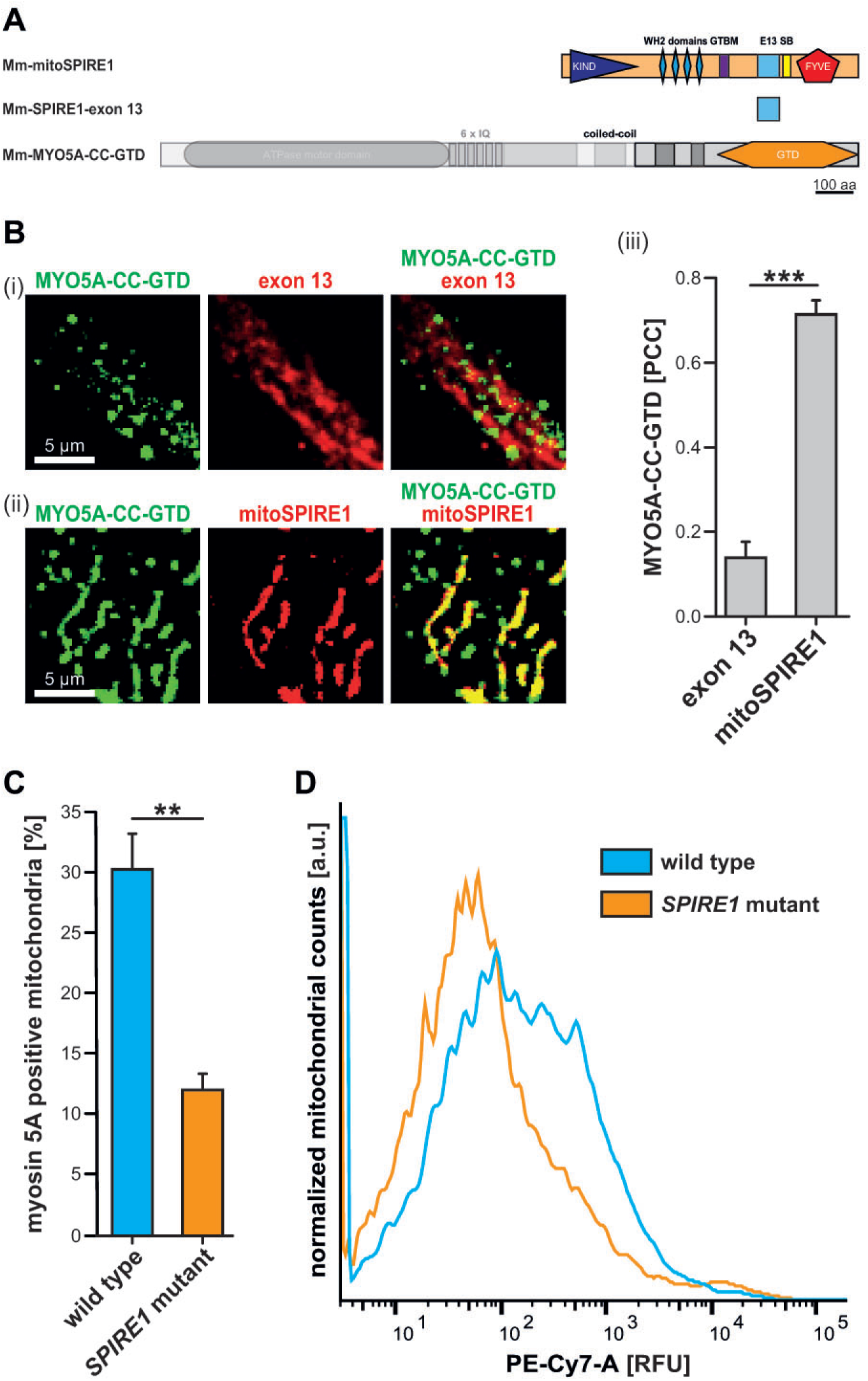
mitoSPIRE1 targets MYO5A proteins towards mitochondrial membranes. **(A)** Schematic overview on proteins used in the colocalization experiment. For a better overview the fluorescence tags (mStrawberry or GFP) are not shown. **(B)** mitoSPIRE1, isolated exon 13 and C-terminal MYO5A proteins were tagged with fluorescent proteins, transiently co-expressed in HeLa cells and analysed by fluorescence microscopy. **(i)** mStrawberry tagged isolated exon 13 sequences (exon 13, *red*) localize to mitochondria. In contrast eGFP tagged C-terminal MYO5A proteins containing coiled-coil regions and the globular tail domain (MYO5A-CC-GTD, *green*) show a typical vesicular localization which is distinct from mitochondria. **(ii)** Co-expression of mStrawberry tagged full-length mitoSPIRE1 (mitoSPIRE1, *red*) and MYO5A-CC-GTD (*green*) proteins shows mitochondrial localization of mitoSPIRE1 and targeting of MYO5A-CC-GTD to mitochondrial membranes in contrast to (i). **(iii)** The extend of colocalization of MYO5A-CC-GTD with isolated exon 13 and full-length mitoSPIRE1 proteins, respectively, was further quantified by determining their Pearson’s correlation coefficients (PCC) as shown in a bar diagram. Each bar represents the mean PCC values for each co-expression, *error bars* indicate SEM. At least 6 cells were analyzed per condition. T-test was performed as post-hoc analysis: *** = p < 0.001. **(C)** The percentage of myosin 5A positive mitochondria was determined in wild-type and *SPIRE1* mutant immortalized mouse embryonic fibroblasts (iMEFs) by fluorescence-activated mitochondria sorting (FAMS). MYO5A proteins at the mitochondria surface were stained with specific antibodies which are fluorescently tagged with PE-Cy7. T-test as post-hoc analysis: ** = p < 0.01. **(D)** PE-Cy7 intensity (horizontal axis) against normalized mitochondrial counts (vertical axis) from iMEFs of *SPIRE1* mutant (orange) and wild-type (blue) mice of the FAMS experiment (B (i)) is plotted in a histogram. iMEFs from *SPIRE1* mutant mice have less mitochondria with a high PE-Cy7 fluorescence intensity compared to mitochondria from wild-type iMEFs suggesting a reduced number of mitochondria from *SPIRE1* mutant iMEFs having MYO5A on their surface compared to wild-type mitochondria. Data are normalized and K-S probability > 99.9 %. *RFU*, relative fluorescence units; *a*.*u*., arbitrary units.

**Fig. 7.**
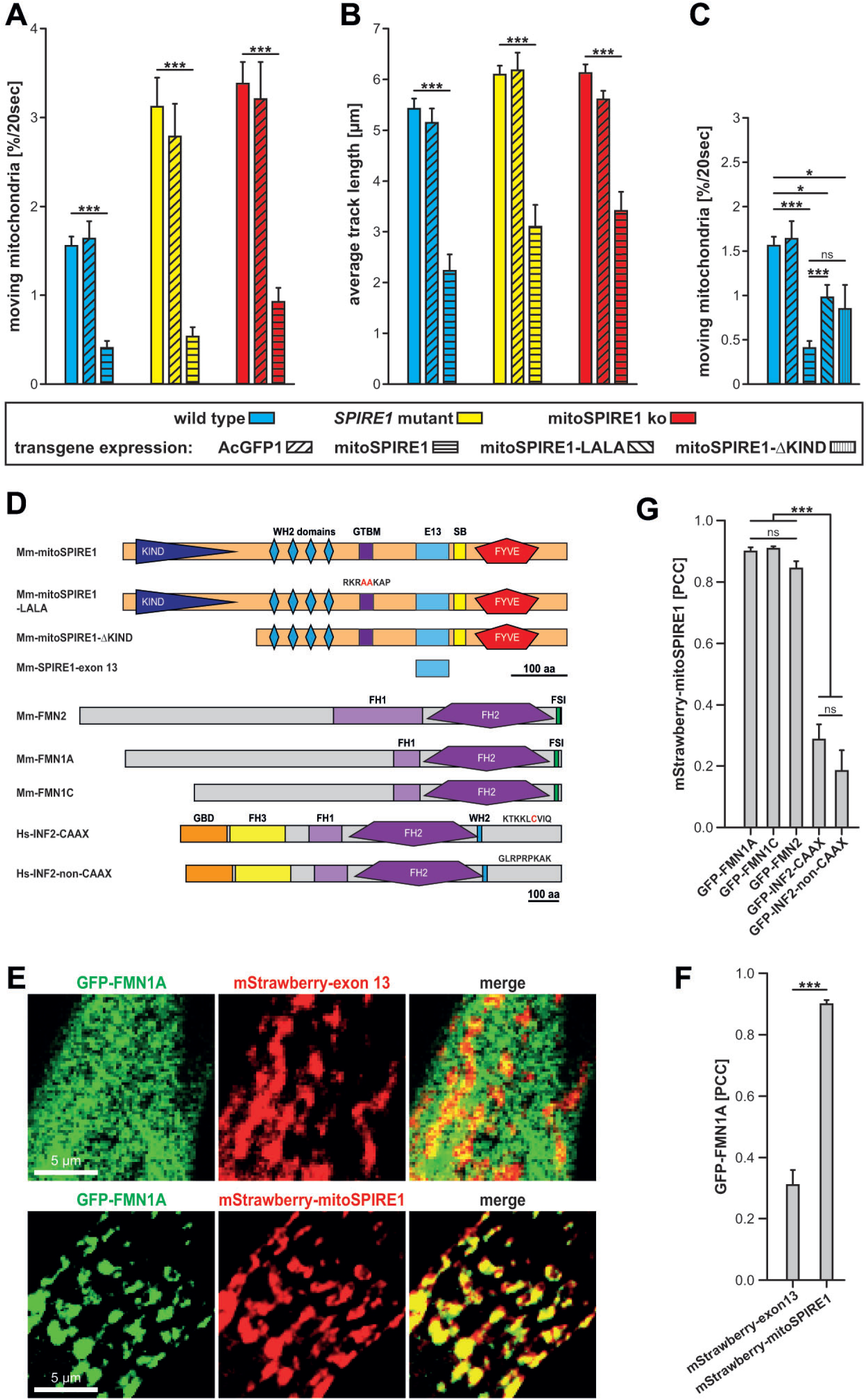
Transgenic overexpression of mitoSPIRE1 stalls mitochondrial motility. Motility of mitochondria was measured in a time range of 20 seconds in primary mouse embryonic fibroblasts (pMEFs) following respective transfections, mitochondria migratory tracks analyzed by IMARIS software. Motility of mitochondria was grouped into three different classifications according to the measured migratory track length: *fixed, wiggle* and *moving* mitochondria. Only *moving* mitochondria were used for further motility analysis. **(A)** The percentage and **(B)** the average track length of *moving* mitochondria is shown in bar diagrams for pMEFs isolated from wild-type (*blue*), *SPIRE1* mutant (*yellow*) and mitoSPIRE1 KO (*red*) mice. For each cell type analysis data are presented for the native state (untransfected control) and overexpression of AcGFP1 or full-length mitoSPIRE1 proteins, respectively (refer to *transgene expression*). Transient overexpression of mitoSPIRE1 reduces all measured parameters significantly, independent from the genotype. Each bar represents mean values, *error bars* represent SEM. n > 12 cells per group. Two-way ANOVA with 1 factor species 3 levels and 1 factor expression 3 levels has been performed; in all cases of expression a significant difference was detected; proofed effect size (η^2^_P_ > 0.9); Tukey-Kramer test has been performed as post-hoc analysis: ** = p < 0.01, *** = p < 0.001. Statistics are only indicated for differences between the native state and overexpression of full-length mitoSPIRE1 proteins. **(C)** Wild-type pMEFs were transfected with mitoSPIRE1-LALA (unable to interact with MYO5) or mitoSPIRE1-ΔKIND (unable to interact with FMN subfamily formins) and the number of moving mitochondria was determined. Cells overexpressing mitoSPIRE1-LALA or mitoSPIRE1-ΔKIND show a significantly reduced number of moving mitochondria compared to untransfected and GFP controls. Each bar represents mean values, *error bars* represent SEM. n > 6 cells per group. One-way ANOVA has been performed; a significant difference between expressions was detected; proofed effect size (η^2^ = 0.302); Mann-Whitney-U-test has been performed as post-hoc analysis: ns = not significant, * = p < 0.05, *** = p < 0.001. **(D)** Schematic overview on proteins used in the mitochondrial motility (A - C) or the colocalization (E - G) experiments. For a better overview the fluorescence tags (mStrawberry or GFP) are not shown. Amino acid labeling at the mitoSPIRE1-LALA protein indicate point mutations in the GTBM (L426,427A; highlighted in red). Amino acid labelling at the C-terminal ends of both INF2 isoforms indicate their specific ending. *KIND*, kinase non-catalytic C-lobe domain; *WH2*, Wiskott-Aldrich-Syndrome protein homology 2; *GTBM*, globular tail domain binding motif; *E13*, exon 13; *SB*, SPIRE-box; *FYVE*, FYVE-type zinc-finger; *FH1*, formin homology domain 1; *FH2*, formin homology domain 2; *FSI*, formin-SPIRE Interaction; *GBD*, GTPase-binding domain; *FH3*, formin homology domain 3; *GFP*, green fluorescent protein; *Hs*, Homo sapiens; *Mm*, Mus musculus; *aa*, amino acids; SPIRE1 accession-number: gi|149321426|ref|NC_000084.5|NC_000084 Mus musculus chromosome 18; accession-numbers of further proteins are listed in Table 2. **(E - G)** Fluorescent protein tagged mitoSPIRE1, isolated SPIRE1-exon 13, FMN1 (FMN1A and FMN1C), FMN2 and INF2 (CAAX and NonCAAX) proteins are transiently overexpressed as well as co-expressed in HeLa cells and analyzed by fluorescence microscopy. **(E)** Deconvoluted images of co-expressed proteins show an overlapping localization of FMN1A (GFP-FMN1A, *green*) with mitoSPIRE1 (mStrawberry-mitoSPIRE1, *red*) but not with isolated exon 13 sequences (mStrawberry-exon 13, *red*) where it rather shows a cytosolic distribution. *Scale bars* represent 5 μm. **(F - G)** The colocalization rate of tagged proteins as described above was quantified for the indicated co-expressions by determining their Pearson’s correlation coefficients (PCC) as shown in a bar diagram. Each bar represents mean PCC values for at least six cells analyzed, *error bars* represent SEM. **(F)** GFP-FMN1A shows a significantly higher correlation for colocalization with mitoSPIRE1 compared to isolated exon 13, which indicates that mitoSPIRE1 translocates GFP-FMN1A from the cytosol towards mitochondria. **(G)** All FMN proteins show a significantly higher PCC value for colocalization with mitoSPIRE1 than INF2 proteins. One-way ANOVA was performed; Tukey-Kramer test was performed as post-hoc analysis: ns = not significant, *** = p < 0.001.

Together, our data show that under transient overexpression conditions, the mitochondrial mitoSPIRE1 protein translocates C-terminal MYO5A proteins towards mitochondria and thereby support a function of mitoSPIRE1 as a mitochondrial adaptor protein for MYO5.

In order to test the targeting of endogenous myosin 5A to mitochondria by mitoSPIRE1, we have analysed and compared the MYO5A abundance on mitochondrial membranes from immortalised wild-type and *SPIRE1* mutant mouse embryonic fibroblasts (iMEFs)(Andritschke et al., 2016). We have employed a nanoscale flow cytometry-based method, which can resolve particles in the size range of mitochondria (fluorescence-activated mitochondria sorting; FAMS)(MacDonald et al., 2019) to analyse the expression of MYO5A on the outer mitochondrial membrane. Following detergent-free lysis of the cells, mitochondria have been stained with the fluorescent dye *MitoTracker Red CMXRos* and a Cy7 fluorescent labeled anti-MYO5A mouse monoclonal antibody. The stained mitochondria were then analysed by FAMS (Fig. 6C, D). The comparison of mitochondria from wild-type iMEFs and those of *SPIRE1* mutant iMEFs, which lack the expression of the mitoSPIRE1 protein, showed a strong difference in MYO5A staining. The mutant mitochondria have lost a significant portion of the population of mitochondria showing high immune reactivity with the MYO5A antibody (Fig. 6C, D). These data indicate that SPIRE1 is a major factor in targeting MYO5 function towards mitochondria on an endogenous level. There was, however, still MYO5A immunoreactivity on mitochondria isolated from *SPIRE1* mutant cells, indicating that there are other MYO5A mitochondria-targeting factors, in addition to mitoSPIRE1.

### mitoSPIRE1 gain of function inhibits mitochondrial motility

Our mitochondria tracking experiments showed that mitochondria of cells lacking mitoSPIRE1 function have an increased motility. Next, we asked whether restoring mitoSPIRE1 function in mitoSPIRE1 knockout and *SPIRE1* mutant cells would reduce the increased motility. We therefore have transiently expressed AcGFP and AcGFP-tagged mitoSPIRE1 in wild-type, mitoSPIRE1 knockout and *SPIRE1* mutant primary fibroblasts. Live cell imaging and tracking of MitoTracker Orange-stained mitochondria revealed that transient overexpression of mitoSPIRE1 in all three cell types strongly inhibited mitochondrial motility as measured by the number of moving mitochondria and the average track length (Fig. 7A, B). Under mitoSPIRE1 overexpression conditions, the mitochondria were nearly static. The transient expression of AcGFP had no significant effect on mitochondrial motility (Fig. 7A, B, C).

In order to address whether the observed inhibition of mitochondrial motility is dependent on the targeting of MYO5 function and/or the generation of actin filaments, we then transiently expressed mitoSPIRE1 mutant proteins in wild-type iMEF cells and analysed mitochondrial motility by live cell fluorescence microscopy (Fig. 7C). Mutants which cannot interact with myosin 5 (mitoSPIRE1 L426A, L427A; mitoSPIRE1-LALA (Pylypenko et al., 2016)) or FMN-subgroup formins (mitoSPIRE1-ΔKIND (Zeth et al., 2011)) were tested.

SPIRE proteins alone cannot generate actin filaments from profilin/actin (Montaville et al., 2014; Quinlan et al., 2005; Quinlan et al., 2007), which is the major pool of polymerizable actin monomers in cells (Kaiser et al., 1999). Only in complex with FMN-subgroup formins, which interact with the N-terminal SPIRE-KIND domain, the SPIRE proteins contribute to the generation of actin filaments (Pfender et al., 2011)(Alzahofi et al., 2020). A SPIRE1-ΔKIND mutant protein, which cannot interact with FMN-subgroup formins, did not rescue SPIRE1 function in actin/myosin driven melanosome transport (Alzahofi et al., 2020). Both mutant mitoSPIRE1 proteins (mitoSPIRE1-LALA and mitoSPIRE1-ΔKIND) inhibited mitochondrial motility much less efficiently than the wild-type protein (Fig. 7C), indicating that the targeting of MYO5 motor proteins and the cooperation with FMN-subgroup proteins in actin filament generation contributed to the stalling of mitochondrial motility.

The reduced inhibitory function of the mitoSPIRE1-ΔKIND mutant led us to speculate that, similar to MYO5, mitoSPIRE1 might target formin proteins to the mitochondrial membrane. We have tested this by transient co-expression of AcGFP1-tagged formin proteins and mStrawberry-mitoSPIRE1 in human HeLa cells (Fig. 7D, E). Next to the two mouse FMN-subgroup proteins, FMN1 and FMN2 (Pechlivanis et al., 2009), the SPIRE1-KIND domain was shown to have a weak interaction with inverted formin-2 (INF2)(Manor et al., 2015), which has two different splice variants: the cytoplasmic INF2-non-CAAX, and the ER-localised INF2-CAAX isoforms (Ramabhadran et al., 2011)Fig. 7D). In fluorescence microscopy co-localisation studies, we observed only a strong co-localisation with mitoSPIRE1 for the FMN-subgroup formins (Fig. 7G). The INF2 formins showed very low, if any, co-localisation with the mitoSPIRE1 protein. In the case of FMN1, we found an even cytoplasmic distribution in the absence of mitoSPIRE1 without mitochondrial localisation, as marked by the co-expression of the isolated exon 13 encoded sequences (mStrawberry-exon 13; Fig. 7E, F). Co-expression with mitoSPIRE1, however, actively targeted FMN1 towards mitochondria, which was similar to the translocalisation of MYO5A (Figs. 5, 7E, F). Based on the mitochondrial motility studies and the colocalisation experiments, we propose a model in which the observed function of mitoSPIRE1 in stalling mitochondria is mediated by the generation of an actin/myosin network at the mitochondrial membrane (Fig. 8).

**Fig. 8.**
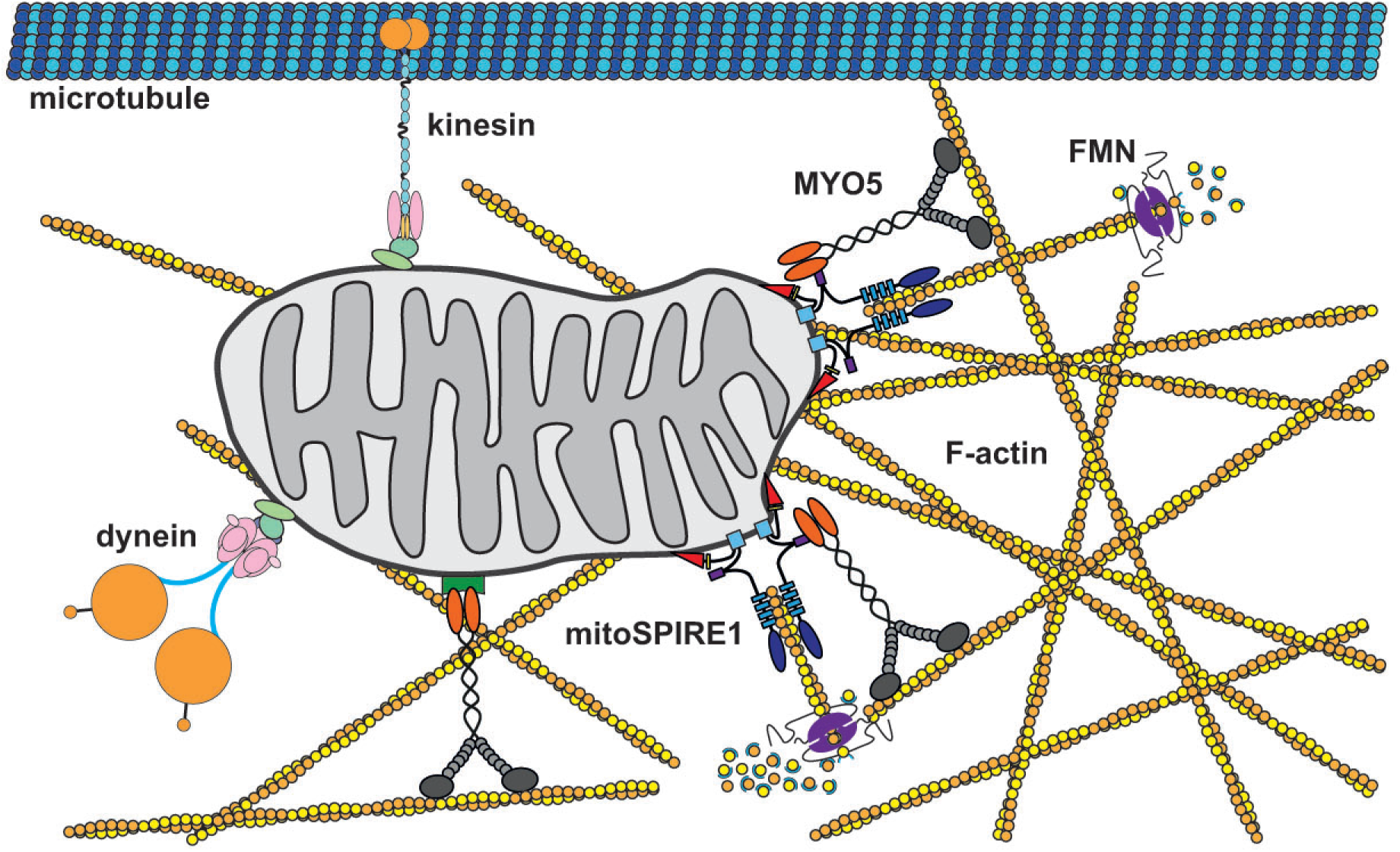
Model of mitoSPIRE1 function in mitochondria anchoring and positioning. Mitochondria are transported along microtubules over long distances via kinesin and dynein motor proteins. In contrast, myosin 5 (MYO5) actin motor proteins might drive short-range local mitochondria transport along actin networks. MYO5 could also influence the processivity of microtubule-based long-range transport of mitochondria. The model suggests that exon 13 drives localization of mitoSPIRE1 to the outer mitochondrial membrane, which facilitates translocation of MYO5 and FMN subfamily formin proteins to mitochondria. Considering the SPIRE-KIND / FMN-FSI interaction a mitochondrion associated mitoSPIRE1 / FMN protein complex might be able to generate actin filaments directly at the mitochondrial surface. Thus, a protein complex, including mitoSPIRE1, FMN subfamily members and MYO5 actin motors, could generate a local mitochondrial actin network, which could ultimately be used by MYO5 to anchor mitochondria at actin filaments, which finally influences mitochondrial motility.

## Discussion

Directional vesicle and organelle transport define the polarisation and structural differentiation of cells throughout evolution. This includes the subcellular localisation of mitochondria, for spatial and temporal organisation of cellular energy demands. Here, we have shown that the mitochondria-bound actin nucleation factor mitoSPIRE1 targets myosin 5 actin motor proteins and FMN-subgroup actin filament generation factors towards mitochondria, and thereby negatively regulates mitochondrial motility. Our data indicate that mitoSPIRE1 acts in mitochondrial positioning and anchoring rather than in active transport processes (Fig. 8). This is in contrast to vesicular SPIRE functions where SPIRE proteins cooperate with FMN-subgroup formins in the generation of actin filaments, which serve as transport tracks for processive myosin 5 motors.

Among the myosin motor protein family, the animal class 5 myosins and their yeast homologs (Myo2p, Myo4p) have a distinguished function in cargo transport (Hammer and Sellers, 2011). The specific ATP/ADP cycle of the class 5 myosin motor domain, which mediates the interaction with actin filaments, enables individual dimeric motors to perform a hand-over-hand processive walk; traveling many steps without dissociating from actin filaments. Mammalian genomes encode three different class 5 myosins: MYO5A, MYO5B and MYO5C (Hammer and Sellers, 2011; Kollmar and Muhlhausen, 2017). All three motors have been shown *in vitro* to processively walk along actin filaments (Heissler et al., 2017; Mehta et al., 1999; Sakamoto et al., 2008; Sladewski et al., 2016). MYO5A and MYO5B motors can walk on single actin filaments, whereas MYO5C requires actin filament bundles for processivity (Sladewski et al., 2016). Cellular long-range transport processes have been shown to depend on MYO5B (RAB11 vesicles)(Schuh, 2011) and MYO5A (RAB27 melanosomes)(Alzahofi et al., 2020) motor activity, where both motors transport cargoes on SPIRE/formin-generated actin meshworks. MYO5C functions in secretory transport processes, and it has been suggested that MYO5C transports exocrine secretory vesicles to the apical membrane on actin filament bundles generated by the formin mDIA1 (Geron et al., 2013).

Next, with regard to active transport functions, mammalian MYO5 has been reported to stall microtubule-based transport and mediate vesicle anchoring. The function of MYO5A in neuronal axonal sorting has been well characterized, whereby MYO5A contributes to stopping the transport of vesicles carrying dendritic cargo proteins in the actin filament-rich axonal initial segment (Janssen et al., 2017; Lewis et al., 2009; Watanabe et al., 2012). By employing a transgenic chemically induced heterodimerisation system, it has been shown that kinesin driven vesicles were arrested in the axon initial segment upon MYO5B recruitment and accumulated in actin-rich hotspots (Janssen et al., 2017). Further evidence for an anchoring function of MYO5A came very recently from a study on dendritic vesicle transport, where MYO5A was found to act as an active tether— mediating long-term stalling of lysosomes at actin patches located at the base of dendritic spines (van Bommel et al., 2019). The MYO5A lysosome anchoring function may resemble the inhibitory opposing role of *Drosophila* MYO5 in axonal mitochondrial transport (Pathak et al., 2010).

It is unknown today what differs between motility and stalling functions of MYO5 motor proteins. The number of motor proteins on an individual vesicle or organelle could be the determining factor. For MYO5A-driven melanosome transport, theoretical considerations have suggested that there are only one or two motor proteins active per cargo (Snider et al., 2004). The motor proteins can also be considered as ADP/ATP-regulated actin filament-binding proteins. This would imply that a higher number of active motors, in the absence of a synchronous ADP/ATP cycle, most likely would fix the vesicles to the actin filaments rather than moving them. In this respect, it would be interesting to quantify and compare the number of SPIRE and MYO5 active motor proteins on melanosomes and mitochondria in order to address the difference of SPIRE/MYO5 function in vesicle motility and mitochondria transport stalling.

In addition to myosin motor protein targeting, our results show that mitoSPIRE1 also recruits the formin actin filament nucleation and elongation factor, FMN1, to the outer mitochondrial membrane. The mitoSPIRE1-KIND domain, which mediates the interaction of SPIRE1 with FMN-subgroup formins (Pechlivanis et al., 2009) contributes to the mitochondrial motility inhibitory effect of mitoSPIRE1 (Fig. 7). This suggests that mitoSPIRE1, in cooperation with formins, could generate actin patches around mitochondria which stall mitochondrial motility. The regulation of mitoSPIRE1 interaction with mitochondria by MYO5 and FMN would provide a mechanism to position and anchor mitochondria at distinct subcellular regions independent of already existing structures of the actin cytoskeleton and in response to signalling cues. In contrast to MYO5, which is regulated by calcium signalling, RAB GTPases, SPIRE, and melanophilin proteins (Hammer and Sellers, 2011; Welz and Kerkhoff, 2019), the regulation of mitoSPIRE1 and FMN-subgroup formins, as well as their integration into signal transduction cascades is virtually not understood today.

In vesicle transport processes, SPIRE proteins are targeted to vesicle membranes by direct interaction with RAB GTPases and/or the interaction of the modified FYVE (mFYVE) zinc finger with the vesicle membrane (Alzahofi et al., 2020; Kollmar et al., 2019; (Tittel et al., 2015). SPIRE is thought to open up from an autoinhibitory N-terminal KIND and C-terminal mFYVE domain interaction, upon the interaction of the mFYVE zinc finger with membranes (Tittel et al., 2015). The release of the KIND/mFYVE interaction subsequently enables the interaction of the KIND domain with FMN-subgroup formins. MitoSPIRE1 could also form an autoinhibited conformation by the KIND/mFYVE interaction. As an initial step towards addressing mitoSPIRE1 regulation, it would be interesting to analyse whether the KIND/mFYVE interaction is inhibited upon the interaction of mitoSPIRE1 with the outer mitochondrial membrane. Our data indicate that both domains are located in the cytoplasm, which would allow a potential auto-inhibitory interaction if the mFYVE zinc finger is not interacting with the mitochondrial membrane.

The novel mitoSPIRE1 knockout mouse generated here were viable and are currently phenotypically analysed. Behavioural studies revealed that the *SPIRE1* mutant mouse, which lacks vesicular and mitochondrial SPIRE1 functions had an increase in fear (Pleiser et al., 2014). Recent findings show that actin-dependent positioning of mitochondria is important for morphological synaptic plasticity (Rangaraju et al., 2019), suggesting a possible role of mitoSPIRE1 in neuronal anchoring of mitochondria during learning and memory, which may influence fear learning, memory and expression. Future behavioural studies comparing *SPIRE1* mutant mice and mitoSPIRE1 knockout mice will reveal if the mitochondrial SPIRE1 function contributes to the fear phenotype.

We found the highest expression of *mitoSPIRE1* in the kidney. The filtration functions of cells lining the kidney tubular system are energetically demanding and require excessive amounts of ATP generated by mitochondria (Bhargava and Schnellmann, 2017). Several cell types within the kidney tubular system have a polarised localisation of mitochondria, with a high enrichment at the basolateral (blood stream) side (Bhargava and Schnellmann, 2017; Subramanya and Ellison, 2014). It will be interesting to analyse whether mitoSPIRE1 is involved in the localisation and anchoring of mitochondria in polarised kidney cells.

Our study presented here provides new insight into the function of SPIRE actin/myosin organisers and exposes a novel approach to address the role of the actin cytoskeleton in the exciting cell biology of mitochondrial dynamics.

## Materials and Methods

### Identification of alternatively spliced exons in *SPIRE* genes

*SPIRE* genes have been identified and annotated previously (Kollmar et al., 2019). All sequences are available at CyMoBase (www.cymobase.org)(Odronitz and Kollmar, 2006). Based on these sequences, available EST and cDNA data was searched for putative alternatively spliced exons. This was done by performing TBLASTN searches using most of the SPIRE protein sequences as query against GenBank nucleotide collection and expressed sequence tags databases. Identified alternatively spliced exons were verified by localisation in the respective genomic DNA sequences. Because cDNA/EST data are incomplete and only available for a few species, the respective genomic regions (intronic regions where the alternatively spliced exons were expected) of the *SPIRE* genes of related species were manually searched for sequences homologous to the identified alternatively spliced exons. Absence of the alternatively spliced exon 13 in fish *SPIRE1B* genes is evident from the short introns in the respective genes, which would easily allow identification of a homologous exon if such an exon were present. In addition, absence is confirmed by analysing multiple species of the same branch. The SPIRE sequence alignment is available from CyMoBase or upon request from the authors. Gene structures were reconstructed using WebScipio (Hatje et al., 2013). The illustrated cartoon (Figure 2A) presents the multiple protein sequence alignment of selected SPIRE1 homologs showing the regions of and around the alternatively spliced exon 13.

### Prediction of secondary protein structure

The online tool Protein Homology / analogY Recognition Engine V 2.0 (PHYRE, intense mode; Structural Bioinformatics Group, London, UK)(Kelley et al., 2015) was used to predict the secondary protein structure of the alternatively spliced exon 13 from the *SPIRE1* gene and the resulting raw structure was further processed in PyMOL (Schrödinger LLC, New York, USA).

### Primer design

Primer pairs for polymerase chain reaction (PCR) and quantitative real-time PCR (qPCR) to amplify specific genomic DNA and cDNA sequences were designed using the web-based tool Primer3 (http://primer3.ut.ee)(Untergasser et al., 2012). To rule out unspecific binding events, the sequence specificity of each primer pair was checked by performing a Primer-BLAST search against the genomic DNA database (www.ncbi.nlm.nih.gov/blast). Primer oligonucleotides consist of 18 to 30 nucleotides and have a GC content of 40 - 60 %. qPCR primer pairs were chosen to amplify exon specific sequences with lengths less than 150 base pairs and flank a region that contains at least one intron. PCR primer pairs were chosen to amplify fragment lengths less than 1,000 base pairs. All primers were synthesized by Sigma-Aldrich (St. Louis, USA) and purified by HPLC. Lyophilized primers were resuspended in DEPC-H_2_O and stored at −20 °C. Prior to use in qPCR, all primers were tested for their capacity to amplify a specific band in PCR and checked by agarose gel separation.

### Qualitative PCR

Standard qualitative PCR amplification of genomic DNA or cDNA fragments was performed to check for abundance of specific nucleotide sequences. Total RNA from cells of interest was isolated using the NucleoSpin RNA Kit from Macherey-Nagel (Düren, Germany) according to manufacturer’s recommendation. Amount and purity of isolated total RNA was measured in spectrophotometric analysis by usage of the Nanophotometer^®^ (Implen GmbH, Munich, Germany). Isolated total RNA was used as a template for cDNA generation by reverse transcription employing the QuantiNova Reverse Transcription Kit (Qiagen, Hilden, Germany) as recommended by Qiagen. Qualitative PCR was set up in 200 μl PCR-SoftTubes (Biozym Scientific GmbH, Hessisch Oldendorf, Germany) using Q5 High-Fidelity DNA Polymerase (New England Biolabs (NEB), Frankfurt am Main, Germany) according to manufacturer’s recommendation. The specific length of PCR products was analysed by agarose gel electrophoresis. The primers used are shown in Table 1.

### Quantitative real-time PCR

Male C57BL/6N mice were ordered from Charles River Laboratories (Sulzfeld, Germany) at an age of 6 weeks. Immediately after arrival mice were sacrificed by cervical dislocation and organs of interest were isolated. Organs were weighed and stored in an adequate volume of *RNA later* (Qiagen) at 4 °C, which inhibits RNA degradation. For total RNA preparations of organs, the RNeasy Midi Kit (Qiagen) was used according to manufacturer’s recommendation. Disruption and homogenization of organs was done with the TissueRuptor (Qiagen). In order to determine amount and purity of isolated total RNA, a spectrophotometric analysis was done as described above. Sample integrity of isolated RNA was measured with the QIAxcel (Qiagen) using the QIAxcel RNA QC Kit v2.0 (Qiagen) according to protocol. Subsequently cDNA was generated employing the QuantiNova Reverse Transcription Kit (Qiagen) according to recommendations. In this method, the ability of the enzyme reverse transcriptase enables the synthesis of DNA from an RNA template. The reverse transcription kit includes a genomic DNA removal step. An internal control RNA was employed to verify successful reverse transcription. Absolute quantification by quantitative real-time PCR (qPCR) was performed with the Rotor-Gene Q thermocycler (Qiagen) and the QuantiNova SYBR Green PCR Kit. Each reaction was set up in triplicates and C_T_ values were calculated by the Qiagen Rotor-Gene Q Series Software. For absolute quantification of sample mRNA copy numbers expression vectors encoding mouse *SPIRE1, SPIRE1-E9, mitoSPIRE1* and *SPIRE2* cDNAs were used to calculate a standard curve with known copy number concentrations. Therefore, plasmid DNA was linearized by restriction digest and serially diluted in water. The copy number of standard DNA molecules was determined by the following formula (from the script “Critical Factors for

Successful Real Time PCR”, Qiagen):

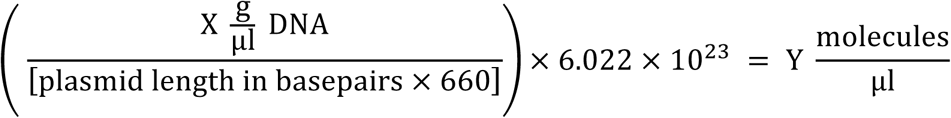

CT values were compared to those of the standard curve with known concentrations to calculate the sample mRNA copy numbers. The primer pairs used for quantitative PCRs are shown in Table 1. The primer pair for “*SPIRE1*” amplifies a fragment present in all three SPIRE1 splice variants whereas the primer pairs for “*SPIRE1-E9*” and “*mitoSPIRE1*” are specific for the distinct splice variant. To determine the exact copy number of *SPIRE1* mRNA, the calculated copy numbers of *SPIRE1-E9* and *mitoSPIRE1* were subtracted from the calculated total *SPIRE1* copy number.

### Cloning of bacterial and mammalian protein expression vectors

Expression vectors were generated by standard cloning techniques using AccuPrime Pfx DNA polymerase (Thermo Fisher, Waltham, MA, USA), restriction endonucleases and T4 DNA ligase (both from NEB). Point mutants were generated by site-directed mutagenesis using the In-Fusion HD cloning kit (TakaraBio/Clontech, Saint-Germain-en-Laye, France). Sequence correctness was verified by sequencing (LGC Genomics, Berlin, Germany). Table 2 shows details of vectors used in this study.

### Cell culture

HEK293, HeLa cells (both from ATCC, Manassas, Virginia, USA) and primary mouse embryonic fibroblasts (pMEFs) were cultured in *Full Medium* (Dulbecco’s Modified Eagle’s Medium (DMEM; Thermo Fisher) supplemented with 10% (v/v) fetal calf serum (FCSIII; GE Healthcare Life Sciences, HyClone), 2 mM L-glutamine (Thermo Fisher), penicillin (100 units/ml; Thermo Fisher) and streptomycin (100 μg/ml; Thermo Fisher)) at 37 °C, 5 % CO_2_, 95% humidity and were passaged regularly at 80% confluency. Transfections with plasmid DNA were performed using Lipofectamine2000 reagent (Thermo Fisher) according to manufacturer’s recommendation.

### Colocalization studies

The extent of colocalization of fluorescently tagged proteins or immunostained proteins was analyzed using the ImageJ (V2.0.0) plug-in Coloc2 (National Institute of Health, Bethesda, USA). Here, the colocalization rate is indicated by the Pearson’s Correlation Coefficient (PCC) as a statistical measure to unravel a linear correlation between the intensity of different fluorescent signals. A PCC value of 1 indicates a perfect colocalization, 0 indicates a random colocalization and a PCC value of −1 indicates a mutually exclusive localization of the analyzed signals. To take the noise of each image into account and to gain an objective evaluation of the PCC significance, a Costes significance test was performed. To do so, the pixels in one image were scrambled randomly and the correlation with the other (unscrambled) image was measured. Significance regarding correlation was observed when at least 95 % of randomized images show a PCC less than that of the original image, meaning that the probability for the measured correlation of two colors is significantly greater than the correlation of random overlap (Costes et al., 2004; Pompey et al., 2013).

### Immunostaining

HeLa cells were seeded on microscope cover glasses and transfected to transiently express fluorescently tagged SPIRE, FMN, INF2 and MYO5 proteins, respectively. Cells were fixed with paraformaldehyde (3.7 % in 1x PBS; Carl Roth, Karlsruhe, Germany) for 20 min at 4°C and subsequently permeabilized using 0.2% Triton X-100 (in 1x PBS; Sigma-Aldrich) for 3.5 min, room temperature. In one experiment 20 μM Digitonin (in DEPC-H_2_O, Sigma-Aldrich) for 3 min, room temperature, was used to permeabilize only the cell membrane. Digitonin is a mild permeabilizer which specifically targets cholesterol-rich membranes such as the plasma membrane, whereas the mitochondria membranes are not affected due to their low concentration of cholesterol. Cells were incubated with anti-cytochrome C (33-8200, 5 μg/ml, mouse monoclonal, Thermo Fisher) and anti-SPIRE1 (SA-2133, 50 μg/ml, rabbit polyclonal; Schumacher et al., 2004) antibodies, respectively, for 1 hour at room temperature, and conjugated anti-mouse TRITC (715-025-151, 9.75 μg/ml, from donkey; Dianova, Hamburg, Germany) and anti-rabbit Cy5 (711-175-152, 9.75 μg/ml, from donkey, Dianova) secondary antibodies, respectively, for 1 h at room temperature avoiding exposure to light. Finally, cells were mounted on microscope slides with Mowiol (Merck, Darmstadt, Germany), dried at room temperature in the dark and stored at 4 °C.

### Fluorescence microscopy of fixed and immunostained cells

Fixed cells were analyzed with a Leica AF6000LX fluorescence microscope, equipped with a Leica HCX PL APO 63x/1.3 GLYC objective and a Leica DFC7000 GT digital camera (1,920 × 1,440 pixel (∼ 2.8 MP); 4.54 μm × 4.54 μm pixel size; all from Leica). 3D stacks were recorded and processed with the Leica deconvolution software module. Images were recorded using the Leica LASX software and further processed with Adobe Illustrator.

### Generation of the mitoSPIRE1 knockout mouse

The mitoSPIRE1 knockout mouse was generated employing the CRISPR / Cas9 gene editing technology. Two targeting constructs were designed (pX458-mitoSPIRE1-1 and pX458-mitoSPIRE1-2) that each express both the gRNA (guide RNA) and the Cas9 nuclease protein. The gRNAs targeted Cas9 nuclease to two distinct locations that are located right in front and behind the exon 13 of the *SPIRE1* gene and a DNA sequence of approximately 179 base pairs is deleted. Potential guide RNAs were screened using CRISPOR tool (www.crispor.tefor.net) and two of them were selected according to the following parameters: cutting site proximity to the location of interest, off target score and Doench score (Doench et al., 2016). The analysis resulted in two gRNAs (gRNA1: TAGGAACCTGAACAATGGAA<u>GGG</u> and gRNA2: CAGCAGGTGGAAACCCTCAA<u>AGG</u>; PAM sequence underlined) that were satisfactory to our requirements. Both gRNAs were individually cloned into pX458 plasmid (Addgene, Watertown, USA), as described Ran and coworkers (Ran et al., 2013). F1.129S2; C57BL/6N embryonic stem cells (ES cells) were transfected with the above-mentioned gRNA and donor template, following the protocol described in Yang and coworkers (Yang et al., 2014). 1 × 10^6^ ES cells were plated in two 10 cm petri dishes coated with gelatine and cultured in N2B27+2i+LIF medium. 4 hours later the cell culture was transfected with a mixture of 5 µg pX458-sgRNA1 + 5 µg pX458-sgRNA2 + 15 µl FuGene following the vendor’s instructions. 48 h later, GFP-positive cells were sorted by FACS and re-plated in gelatine-coated 10 cm Petri dishes (25,000 cells / dish). After one week in culture, individual ES cell colonies were picked manually, disaggregated and plated on a feeder cell layer in 96-well plates. This clone set was split in triplicates: two of the copies were frozen and the third was expanded up to confluent 24-well plate wells for DNA extraction and genotyping. ESC clones were genotyped by PCR reaction using the primers of Table 1 (*mitoSPIRE1* (PCR, genotyping)). Clones that were validated for a homozygous knockout by sequencing were further processed. Once the ESC clones carrying the deletion were identified by PCR, one of the two frozen clone-set replicas were thawed and the selected clones were put in culture on a feeder cell layer in 96-well plates with N2B27+2i+LIF medium. Subsequently, these clones were then expanded in the same culture conditions transferred to 6-well plate wells. Cells were disaggregated to single cell suspension and microinjected into morula stage C57BL6/N embryos, at a ratio of 3 - 5 cells per morula. The injected embryos were transferred to pseudo-pregnant recipient CD1 females to be carried to term.

### Recombinant protein expression and purification

Recombinant GST-mm-FMN2-eFSI (GST-FMN2-eFSI) proteins were expressed in *Escherichia coli* Rosetta bacterial cells (Merck Millipore). Bacteria were cultured in LB medium (100 mg/l ampicillin, 34 mg/l chloramphenicol) at 37°C until an OD_600nm_ of 0.6 - 0.8. Protein expression was induced by 0.2 mM Isopropyl-ß-D-thiogalactopyranoside (IPTG; Sigma-Aldrich) and continued at 20°C for 20 h. Bacteria were harvested and lysed by ultra-sonication. Soluble proteins were purified in two consecutive steps. GSH-Sepharose 4B beads (GE Healthcare Life Sciences) were used first for an affinity-based batch purification process which was followed by size exclusion chromatography using an ÄKTApurifier system and a High Load 16/60 Superdex 200 SEC column (both GE Healthcare Life Sciences). Proteins were concentrated by ultrafiltration using Amicon Ultra centrifugal filters (Merck Millipore) with 50 kDa cut off. The final protein purity was estimated by SDS-PAGE and Coomassie staining.

### GST-pulldown from wild-type and mitoSPIRE1 knock out fibroblast lysates

For GST-pulldowns eight 10 cm cell culture dishes with 80% confluent primary mouse embryonic fibroblasts (pMEFs) from both, wild-type and mitoSPIRE1 knock out mice, were used. Cells were lysed in lysis buffer (25 mM Tris-HCl pH 7.4, 150 mM NaCl, 5 mM MgCl_2_, 10% (v/v) glycerol, 0.1% (v/v) Nonidet P-40, 1 mM PMSF, protease inhibitor cocktail) with 1 ml buffer was used for cells from 4 dishes, and afterwards centrifuged at 20,000 × g, 4 °C, 20 min to remove insoluble debris. For subsequent GST-pulldown assays 40 µg GST-FMN2-eFSI and 20 µg GST protein as control were coupled to GSH-Sepharose 4B beads for 1 h, 4°C on a rotating wheel followed by washing twice with pulldown buffer (25 mM Tris-HCl pH 7.4, 150 mM NaCl, 5 mM MgCl_2_, 10 % (v/v) glycerol, 0.1 % (v/v) Nonidet P-40). Cleared lysates from each cell line were pooled to get equal protein concentrations throughout all pulldown samples and a volume of 1 ml was subsequently incubated with the coupled beads for 2 h at 4 °C on a rotating wheel. Beads were washed four times with pulldown buffer and bound proteins were eluted with 1x Laemmli buffer, denatured at 95 °C for 10 min and then analyzed by immunoblotting using anti-SPIRE1 (SA-2133, rabbit polyclonal, 0.5 µg/ml; Schumacher et al. 2004) primary antibody and horseradish peroxidase linked anti-rabbit IgG secondary antibody (1:5000, from donkey, GE Healthcare Life Sciences). Signal was detected by chemiluminescence (Luminata Forte Western HRP substrate; Merck Millipore) and recorded with an Image Quant LAS4000 system (GE Healthcare Life Sciences).

### Isolation of pMEFs from mouse embryos

For successful isolation of primary mouse embryonic fibroblasts (pMEFs) from homozygous mitoSPIRE1 knockout and wild-type embryos heterozygous mitoSPIRE1 knockout mice were crossbred. For further experiments isolated pMEFs were only considered from litters containing all three genotypes to reduce litter specific variances. At embryonic day 14.5 pregnant mice were killed by cervical dislocation. The uterus was removed, embryos were isolated and stored in a petri dish with cold sterile 1x PBS. Two limbs of each embryo were collected and used for genotyping of the embryos (see below). Each embryo was handled individually and within 60 minutes after removal of the uterus. After removing all red organs from the embryo, the residual tissue was washed with 1x PBS to remove remaining blood. In the next step the embryo was sliced into fine pieces with two scalpels (Megro, Wesel, Germany) and transferred into a 15 ml conical tube containing 3 ml of ice-cold 0.05 % Trypsin-EDTA solution (Thermo Fisher). Tissues were stored over night at 4°C to allow the Trypsin-EDTA solution to diffuse into it. After each pMEF isolation from different embryos scissors (Megro), forceps (Dumont, Montignez, Switzerland) and scalpels were washed with water and 70 % ethanol to avoid cross-contamination between different genotypes. At the next day the Trypsin-EDTA solution supernatant was discarded and the embryonal tissue was incubated for 15 min at 37 °C in a water bath. 3 ml *Full Medium* (refer to cell culture) was added to each embryo to inactivate Trypsin-EDTA and to facilitate a gentle homogenization of tissue clumps by pipetting gently up and down with a 1000 μl pipette tip (Axygen, Corning, USA) and a 200 μl pipette tip (Sarstedt, Nümbrecht, Germany). The cell suspension of each embryo contained pMEFs and was finally splitted on four 10 cm tissue culture plates each containing 10 ml of *Full Medium*. pMEFs were cultured at 37°C, 5% CO_2_, 95% humidity and were passaged regularly at 80% confluency. pMEFs were frozen in liquid nitrogen for long-term storage and thawed as required for experiments. All experiments with pMEFs were done with cells not older than passage five, because proliferation rate of pMEFs in culture decreases dramatically after five passages.

### Genotyping of embryos used for pMEF isolation

Two limbs of each embryo were collected in a 1.5 ml tube and quickly covered with tail lysis buffer (50 mM Tris-EDTA pH 8.0, 0.5 % (w/v) SDS) freshly supplemented with Proteinase K (Carl Roth; final concentration 1 mg/ml). Limb tissue was lysed overnight (∼20 h) at 56 °C. Samples were centrifuged at 20,000 × g at room temperature and supernatant was diluted 1:50 in DEPC-H_2_O. Diluted genomic DNA was utilized for PCR analysis using Taq DNA polymerase (NEB) and genotyping primers indicated in Table 1. In detail, for mitoSPIRE1 knockout genotyping we used a primer pair starting in the *SPIRE1 intron 12* and ending in the belonging *intron 13* sequence to confirm the absence of genomic *SPIRE1 exon 13*. If *exon 13* is included in the *SPIRE1* gene a band of 749 base pairs is generated. In contrast, the specific primer pairs generate a 575 base pair DNA fragment when *exon 13* is absent. For genotyping of *SPIRE1* mutant mice two primer pairs were used. In this case, wild-type samples show a specific band of 269 base pairs when ‘Primer pair 1: wild-type / SPIRE1 mutant’ was used, while *SPIRE1* mutant samples did not show a specific band. In contrast, *SPIRE1* mutant samples showed a 142 base pair DNA fragment using ‘Primer pair 2: wild-type / SPIRE1 mutant’, while wild-type samples did not show a specific PCR product. Specific bands were visualized by agarose gel electrophoresis.

### Mitochondrial staining for fluorescence live cell imaging

In order to fluorescently label mitochondrial membranes in living HeLa cells the cell permeable compound MitoTracker Orange (Thermo Fisher) was used. HeLa cells were seeded in glass-bottom tissue culture dishes (WillCo-Dish, 40 mm; WillCo Wells B.V., Amsterdam, The Netherlands) 20 h prior to the experiment. On the next day cells were washed in Optimem (Thermo Fisher) and incubated with 2 ml of 5 nM MitoTracker Orange (diluted in Optimem) for 20 min at 37 °C, 5 % CO_2_. Cells were washed again with Optimem to get rid of excess staining solution and covered with Optimem for subsequent fluorescence live cell imaging.

### Quantification of mitochondrial motility

Mitochondrial motility of primary mouse embryonic fibroblasts (pMEFs) from wild-type, *SPIRE1* mutant and mitoSPIRE1 knockout mice was evaluated with IMARIS (Bitplane, Zürich, Switzerland). First mitochondria were stained with MitoTracker Orange as described above and recorded by live cell fluorescence imaging with a Leica AM TIRF MC microscope equipped with a Leica HCX PL APO 100x/1.47 oil objective (all from Leica) and a Hamamatsu EM-CCD C9100-02 digital camera (1000 × 1000 pixel (∼ 4.2 MP); 8 μm × 8 μm pixel size; Hamamatsu Photonics, Herrsching am Ammersee, Germany). The data presented here were obtained only by epi-fluorescence imaging. All cells were picked randomly and mitochondria movement from these cells was recorded for 20 seconds (1 image / 500 ms). Movies of motile mitochondria were transferred as Leica project files to IMARIS and movies were processed with the integrated Matlab background subtraction (The MathWorks, Natick, USA). In the following step mitochondria were tracked with the surface tool of IMARIS (Max Distance: 1.5 μm; Max Gap Size: 3 μm). Mitochondria movement was difficult to track automatically, because mitochondria crossed each other and sometimes moved fast leading to loss of tracks or generation of incorrect tracks. For this reason, automatically tracked mitochondria by the surface tool of IMARIS were checked manually and in case of missing or wrong tracks, tracks were corrected manually. Based on the measured tracks IMARIS calculated parameters such as track length, maximal track length or maximal velocity of mitochondria and these results were exported and saved as Excel-files. Data were used to determine the percentage of moving, wiggling and stationary mitochondria in pMEFs from wild-type, *SPIRE1* mutant and mitoSPIRE1 knockout mice. Here, moving mitochondria were determined as mitochondria with a track length > 4 μm / 20 s, while wiggling mitochondria had a track length of 1.5 - 4 μm / 20 s, and stationary mitochondria moved less than 1.5 μm/20 s. The calculated parameters from IMARIS were defined as follows. ‘Track length’ describes the covered distance of moving mitochondria per time interval. ‘Maximal track length’ defines the longest covered distance by one moving mitochondrion of a cell per time interval. ‘Maximal velocity’ determines the highest speed reached by moving mitochondria while transportation. Each parameter was calculated for all moving mitochondria of a cell. Data were summarized in Excel (Microsoft Corporation, Albuquerque, USA) and further processed to generate means of parameters per cell. For the presentation of parameters in a bar diagram, the average of all mean values calculated per cell was evaluated. The same procedure was used to measure mitochondrial motility in case mitoSPIRE1, mitoSPIRE1-LALA or mitoSPIRE1-ΔKIND was transiently overexpressed in pMEFs. Transient overexpression of AcGFP1 was used as a control and all transfections were performed as described above.

### Nanoscale flow cytometry analysis

Cultured cells were detached and collected by centrifugation (300 × g for 5 minutes). The cell pellet was resuspended in FACS buffer (1x PBS pH 7.2, 0.5 % BSA, 2 mM EDTA) at a cell density of 1 × 10^6^ cells/ml. Cells were incubated with 100 nM MitoTracker Red CMXRos (MTR; Thermo Fisher) at 37°C, 5% CO_2_ for 15 minutes. Cell suspensions were pelleted at 300 × g and resuspended in 1 ml of ice-cold hypo-osmotic buffer (RSB Hypo Buffer, Cold Spring Harbor Protocols; 10 mM NaCl, 1.5 mM MgCl_2_, 10 mM Tris-HCl (pH 7.6)). Following a ten-minute swelling period at 4 °C, samples were subjected to 35 strokes of dounce homogenization. Cell lysates were centrifuged at 12,000 × g for 5 minutes at 4 °C. Supernatant was aspirated, and pellets were resuspended in blocking buffer (0.2% BSA and 2% fetal bovine serum in 1x PBS) and incubated at room temperature for 20 minutes. Samples were centrifuged at 12,000 × g for 5 minutes and then incubated with anti-myosin 5A (G-4) antibody (sc-365986, 0.75 μg/ml, mouse monoclonal; Santa Cruz Biotechnology, Dallas, TX, USA) conjugated to PE-Cy7 (ab102903; Abcam, Cambridge, UK). Samples were pelleted at 12,000 × g and resuspended in buffered blood bank saline (0.85% (w/v) isotonic solution; Thermo Fisher) for analysis via nanoscale flow cytometry as previously described (MacDonald et al., 2019). Briefly, assessment of mitochondria was performed using a special ordered BD FACS Aria III, fitted with a PMT detector for forward light scatter (FSC) of a 488 nm laser. Mitochondria were identified by size and positive fluorescence using mitochondrial-specific dye (MTR) in comparison to unstained negative control samples, allowing up to 0.5% false positive events versus controls. Spectral overlap from MTR in PE-Cy7 channel was corrected manually using single color samples. For each sample, 50,000 events in total were analyzed. Data were acquired using BD FACSDiva software (version 8.0.2; Becton Dickinson, Franklin Lakes, USA) and analysis performed using FlowJo (version 10; FlowJo LLC, Ashland, USA).

### Statistical analysis

For all statistical analysis the software SPSS (IBM, Armonk, USA) was used. To compare more than two groups on one level a One-way analysis of variance (ANOVA) was performed, while the comparison of more than two groups on two levels was employed by a Two-way ANOVA. If ANOVA revealed a significant difference and data are distributed normally a Tukey-Kramer test was used as post-hoc analysis, while for data which failed normality test a Mann-Whitney-U-test was performed. Statistical analysis between two groups was performed if normality test was passed by t-test and if normality test failed by Mann-Whitney-U-test. A difference between groups was determined as significant if the alpha level of post-hoc analysis was < 0.05. For groups with n > 25 the effect size was determined as well. In this context, for One-way ANOVA analysis eta squared (η^2^) was calculated to determine the effect size (η^2^ > 0.01 - 0.06, describes a small effect size; η^2^ > 0.06 - 0.14, describes a medium effect size; η^2^ > 0.14 describes a large effect size). For Two-way ANOVA analysis partial eta squared (η^2^_P_) was calculated to determine the effect size (η^2^_P_ > 0.1 - 0.3, describes a small effect size; η^2^_P_ > 0.3 - 0.5, describes a medium effect size; η^2^_P_ > 0.5 describes a large effect size).

## Acknowledgements

We thank Uri Manor for discussion and Roland Wedlich-Söldner for providing us with INF2 expression vectors. We further thank Michael Mahlert for helping to set up the Bitplane Imaris tracking analysis. Felix Straub was a student of a research training group (GRK 2174) of the German Research Foundation (DFG). The work was funded by the prioritiy programme SPP 1464 of the German Research Foundation (KO 2251/13-1 (to MK), KE 447/10-2 (to EK), the individual DFG research grant KE 447/18-1 (to EK) and the National Science Foundation, USA, NSF 1750996 (to DW).

